# Substrate recognition mechanism of the endoplasmic reticulum-associated ubiquitin ligase Doa10

**DOI:** 10.1101/2024.01.09.574907

**Authors:** Kevin Wu, Samuel Itskanov, Diane L. Lynch, Yuanyuan Chen, Aasha Turner, James C. Gumbart, Eunyong Park

**Author notes:** These authors contributed equally to this work (Kevin Wu, Samuel Itskanov).

## Abstract

Doa10 (MARCH6 in metazoans) is a large polytopic membrane-embedded E3 ubiquitin ligase in the endoplasmic reticulum (ER) that plays an important role in quality control of cytosolic and ER proteins. Although Doa10 is highly conserved across eukaryotes, it is not understood how Doa10 recognizes its substrates. Here, we defined the substrate recognition mechanism of Doa10 by structural and functional analyses on *Saccharomyces cerevisiae* Doa10 and its well-defined degron Deg1. Cryo-EM analysis shows that Doa10 has unusual architecture with a large lipid-filled central cavity, and its conserved middle domain forms an additional water-filled lateral tunnel open to the cytosol. Our biochemical data and molecular dynamics simulations suggest that the entrance of the substrate’s degron peptide into the lateral tunnel is required for efficient polyubiquitination. The N- and C-terminal membrane domains of Doa10 seem to form fence-like features to restrict polyubiquitination to those proteins that can access the central cavity and lateral tunnel.

## Introduction

Selective degradation of misfolded and mistargeted proteins constitutes key pathways underpinning cellular protein homeostasis. In eukaryotic cells, the aberrant proteins are marked with polyubiquitin chains by E3 ubiquitin (Ub) ligases and subsequently degraded by the proteasome. The endoplasmic reticulum (ER) serves as a primary site for protein biosynthesis and maturation. Over one third of proteins translocate into the ER lumen or integrate into or associate with the ER membrane, including many proteins that are destined for other organelles ^1–5^. To enable quality control of these proteins, the ER is equipped with a set of membrane-embedded E3 ligases that play central roles in the process known as ER-associated protein degradation (ERAD) ^6–11^. The primary function of these E3 ligases is to recognize and polyubiquitinate aberrant proteins in the ER to enable their selective clearance. Moreover, through coordination with a AAA+ ATPase motor (Cdc48 in yeast and p97 in mammals), ERAD-specific E3 ligases facilitate the removal of proteins from the ER lumen or membrane into the cytosol for proteasomal targeting, a process called retrotranslocation ^12–16^. As the ER forms the most abundant membrane structure as well as a key biosynthetic hub, ERAD constitutes a vital component of protein quality control and regulated proteolysis in eukaryotic cells. Impairment of the ERAD machinery causes ER stress and dysfunction and is implicated in several human diseases ^17,18^. However, the molecular mechanisms by which ERAD-specific E3 ligases mediate the protein quality control processes are incompletely understood.

Conceptually, ERAD can be classified into three categories depending on the topological location of substrate recognition with respect to the ER membrane: (1) the ER lumen, (2) the ER <u>m</u>embrane, and (3) the <u>c</u>ytosol. Respectively, these distinct ERAD pathways are often referred to as ERAD-L, ERAD-M, and ERAD-C ^19–21^. ERAD-L substrates include misfolded lumenal proteins and membrane proteins with a misfolded lumenal domain, whereas ERAD-M substrates include those with misfolding in the intramembrane regions, and ERAD-C substrates are typically proteins with a misfolded cytosolic domain. In addition to misfolding, certain polypeptide segments and posttranslational modification features of substrates also serve as degradation signals (degrons) for ERAD ^22–24^. Generally, the different classes of substrates are recognized by distinct ERAD-specific E3 ligases. In fungal species, most ERAD-L/-M and ERAD-C substrates are handled by Hrd1 and Doa10, respectively, both of which belong to RING-type E3 ligases ^25–27^. In addition to ERAD-C substrates, Doa10 also recognizes certain ERAD-M substrates ^28^. Hrd1 and Doa10 are the two most conserved ERAD-specific E3 ligases across eukaryotic species including humans.

Doa10 (MARCH6/TEB4 in metazoans and SUD1 in plants) is a large multi-spanning membrane protein residing in the ER and inner nuclear membranes ^29^ (Fig. 1a). Its first ∼100 amino acids contain a RING-CH domain that enables its Ub ligase activity through an interaction with E2 Ub-conjugating enzymes. The RING-CH domain is followed by a transmembrane domain (TMD) containing 14 transmembrane segments (TMs), the structure and function of which are poorly characterized. Substrate polyubiquitination by fungal Doa10 requires two E2 proteins Ubc6 and Ubc7 (ref. ^27^). Ubc6 is a tail-anchored (TA) membrane protein in the ER membrane. Ubc7 is a soluble enzyme but localizes to the ER membrane through interaction with the single-spanning ER membrane protein Cue1 (ref. ^30^). Ubc6 is involved in attachment of the first couple of Ub molecules to the substrate, Ubc7 is used for further elongation of the poly-Ub chain ^31^. Doa10 is also shown to interact with Ubx2, a membrane protein that recruits Cdc48 to the ER, facilitating the extraction of substrate polypeptides from the ER membrane into the cytosol in the cases of membrane-associated substrates ^32^. In addition to ER proteins, Doa10 can also polyubiquitinate soluble cytosolic proteins as its substrate recognition mainly occurs on the cytosolic side ^33^.

**Figure 1.**
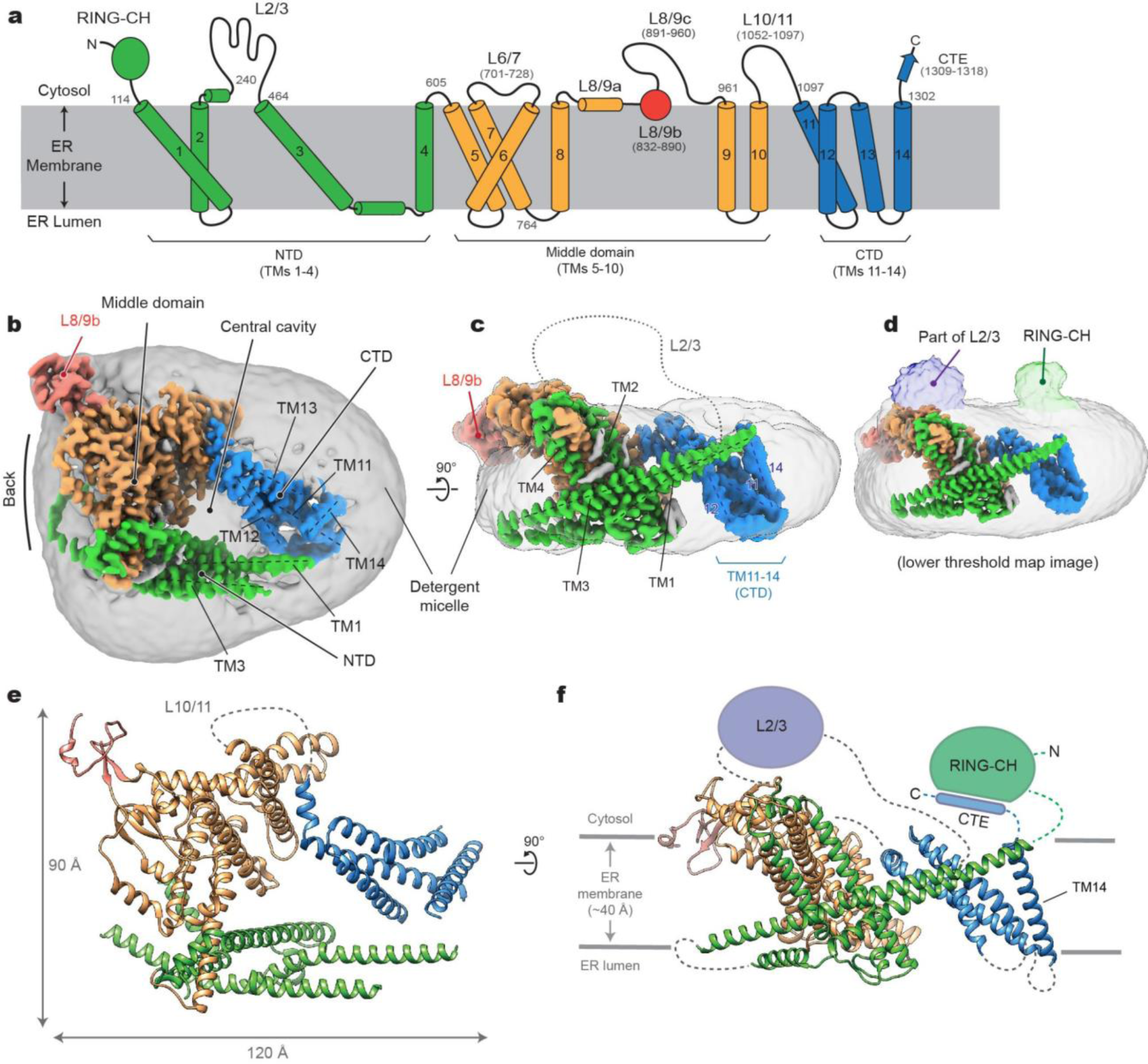
Cryo-EM structure of *S. cerevisiae* Doa10 in detergent micelle. **a**, Schematic diagram of the domain structure of *S. cerevisiae* Doa10. Helices are represented as cylinders (TMs 1-14 are numbered). Numbers in gray indicate amino acid residue number. **b**, Cryo-EM density map of Doa10 viewed from the cytosol. Shown is a high-resolution protein density map overlaid with a lowpass-filtered detergent micelle density (gray). The color scheme is the same as in **a**. **c**, As in **b**, but showing a side view along the membrane plane. **d**, As in **c**, but overlaid a density map with a lower surface threshold to show RING-CH and L2/3 features. **e** and **f**, Atomic model of Doa10 based on the cryo-EM map. Views in **e** and **f** are equivalent to **b** and **c**, respectively. Parts that are unresolved in cryo-EM were schematized.

Currently, the mechanism by which Doa10 recognizes its substrates and coordinates with its E2s for polyubiquitination is unclear. Although a short peptide called Deg1, which was derived from the yeast mating-type protein α2, has been identified as a Doa10-specific degron (Doa10 stands for <u>D</u>egradation <u>o</u>f <u>A</u>lpha2 <u>10</u>) and often used as a model substrate in the studies of Doa10 (ref. ^27,33–35^), it remains unknown how Doa10 recognizes Deg1. Interestingly, certain point mutations in the TMD of Doa10 have been found to alter the turnover rates of Deg1-fused protein substrates, suggesting a role of the TMD of Doa10 in polyubiquitination ^36^. Moreover, it has been suggested that Doa10 itself can also dislocate certain transmembrane substrates from the membrane independently of substrate ubiquitination and Cdc48 (ref. ^37^). Thus, the TMD of Doa10 may possess multiple functions including substrate recognition and retrotranslocation, but the mechanisms underlying these functions remain yet to be understood.

In this study, we present the results of our cryo-electron microscopy (cryo-EM), molecular dynamics (MD), AlphaFold2 modeling, and biochemical analyses of Doa10 from *Saccharomyces cerevisiae*. Our study reveals the highly unusual architecture of Doa10, where its TMD is arranged as a horseshoe-like architecture with its central cavity filled with lipids. The conserved middle domain of Doa10 forms a lateral tunnel-like cavity, the entrance of which faces the central cavity and is open to the cytosol. Mutations in the tunnel impair degradation of Deg1 substrates, and MD simulations suggest that these mutations induce a collapse of the tunnel. Furthermore, we used photocrosslinking to show that the lateral tunnel directly interacts with Deg1 substrates. Using membrane-anchored Deg1 substrates, we further show that a distance of at least 20-30Å between Deg1 and the membrane anchor is required for Deg1 recognition by Doa10. This is likely because fence-like features of the N- and C-terminal portions of TMD restrict membrane proteins from accessing the central cavity and lateral tunnel. Our findings provide important insight into the mechanism by which Doa10 recognizes its substrates. Given the high degree of structural conservation, human MARCH6 is also likely to use the same mechanism to recognize substrates.

## Results

### Cryo-EM analysis of *S. cerevisiae* Doa10

To gain structural insights into the mechanism of Doa10, we first determined a cryo-EM structure of *S. cerevisiae* Doa10. To purify Doa10, we inserted a cleavable green fluorescent protein (GFP) tag at the C-terminus of the chromosomal copy of Doa10 in yeast. We first tested purification of endogenous Doa10 by GFP affinity purification after solubilizing membranes with the lauryl maltose neopentyl glycol (LMNG) detergent. However, the yield and quality of samples were insufficient for structural analysis (Supplementary Fig. 1a, b). We therefore overexpressed Doa10 by replacing its endogenous promoter with a strong galactose-inducible *GAL1* promoter. The purified Doa10 protein in this approach yielded a largely monodisperse peak in size-exclusion chromatography and showed mainly as the full-length band on a SDS gel (Supplementary Fig. 1c, d). Cryo-EM images of the purified Doa10 sample displayed evenly dispersed particles, which produced well defined two-dimensional (2D) class averages (Supplementary Fig. 2a, b).

Single particle cryo-EM analysis of purified Doa10 yielded only one major class, which could be refined to a three-dimensional (3D) reconstruction at 3.2-Å overall resolution (Fig. 1b–d, and Supplementary Figs. 2c and 3 and Table 1). Most of the TMD were well defined, allowing us to build a reliable atomic model (Fig. 1e,f). However, we could not fully register distal parts of the last four TMs (i.e., TMs 11–14) due to their lower local resolution caused by the bending motions of the domain (Supplementary Figs. 3c and 4). We also note that the N-terminal RING-CH domain (residues 1–113) and the loop between TMs 2 and 3 (L2/3; residues 242–463) are only visible at low resolution due to high conformational heterogeneity (Fig. 1d). While we were conducting follow-up biochemical studies, AlphaFold2 was published ^38^. An AlphaFold2 predicted model of yeast Doa10 agrees well with our experimental structure with a root mean square deviation (RMSD) of 2.7 Å over 835 aligned Cɑ atoms (Supplementary Fig. 5a,b). Since our cryo-EM structure cannot model the RING-CH domain and several loops, we also built a hybrid model combining our experimental model and high-confidence regions of the AlphaFold2 model.

### Overall architecture and structural features of Doa10

Doa10 exhibits highly unusual architecture. Overall, the TMD of Doa10 can be divided into the three subregions: the N-terminal domain (NTD), the middle domain, and the C-terminal domain (CTD), which are formed by TMs 1–4, 5–10, and 11–14, respectively (Fig. 1a–c). Viewed from the cytosol, they are arranged in a horseshoe-like fashion (Fig. 1b). Consequently, Doa10 possesses a large enclosed “central” cavity within its TMD. In our cryo-EM map, this cavity is filled with detergent and lipid molecules, suggesting that it would be occupied with lipids in the native membrane. Based on the low-resolution features, the RING-CH domain is placed directly above where TM1 and TM14 join at the tips of the horseshoe-like structure (Fig. 1d).

The TMD of Doa10 is also atypical in that many of its TMs are unusually long and highly tilted (Fig. 1). For example, TM3 is ∼60 amino acids long and tilted by ∼65° from the membrane normal. TMs 1, 5, and 9 are also ∼50 amino acids long and tilted by more than 45°. A few other TMs (TMs 2, 6, and 11) are ∼30–40 amino acids long, substantially longer than lengths of 20–30 amino acids for typical TMs. There are also multiple amphipathic domains. The segment between TMs 3 and 4 includes two amphipathic α-helices lying flat on the lumenal leaflet of the ER membrane. Part of the segment between TMs 8 and 9 (referred to as L8/9b and to be discussed later) forms a globular domain that is partially embedded on the cytosolic leaflet of the ER membrane.

Another atypical feature of Doa10 is a relatively loose packing between its TMs. As a result, the TMD itself contains two sizable intra-TMD cavities each surrounded by TMs 1–5 (cavity 1) and by TMs 5– 10 (cavity 2), respectively, and both are partly continuous from the central cavity (Fig. 2a,b; Supplementary Fig. 3e). Our cryo-EM structure shows that both cavities are occupied by lipids. Interestingly, cavity 2 is occupied with a triglyceride in addition to a phospholipid. The arrangement of TMs 5–10 suggests that these two lipids are unlikely to freely exchange with the bulk lipids, and therefore are likely trapped into the cavity during the folding of Doa10.

**Figure 2.**
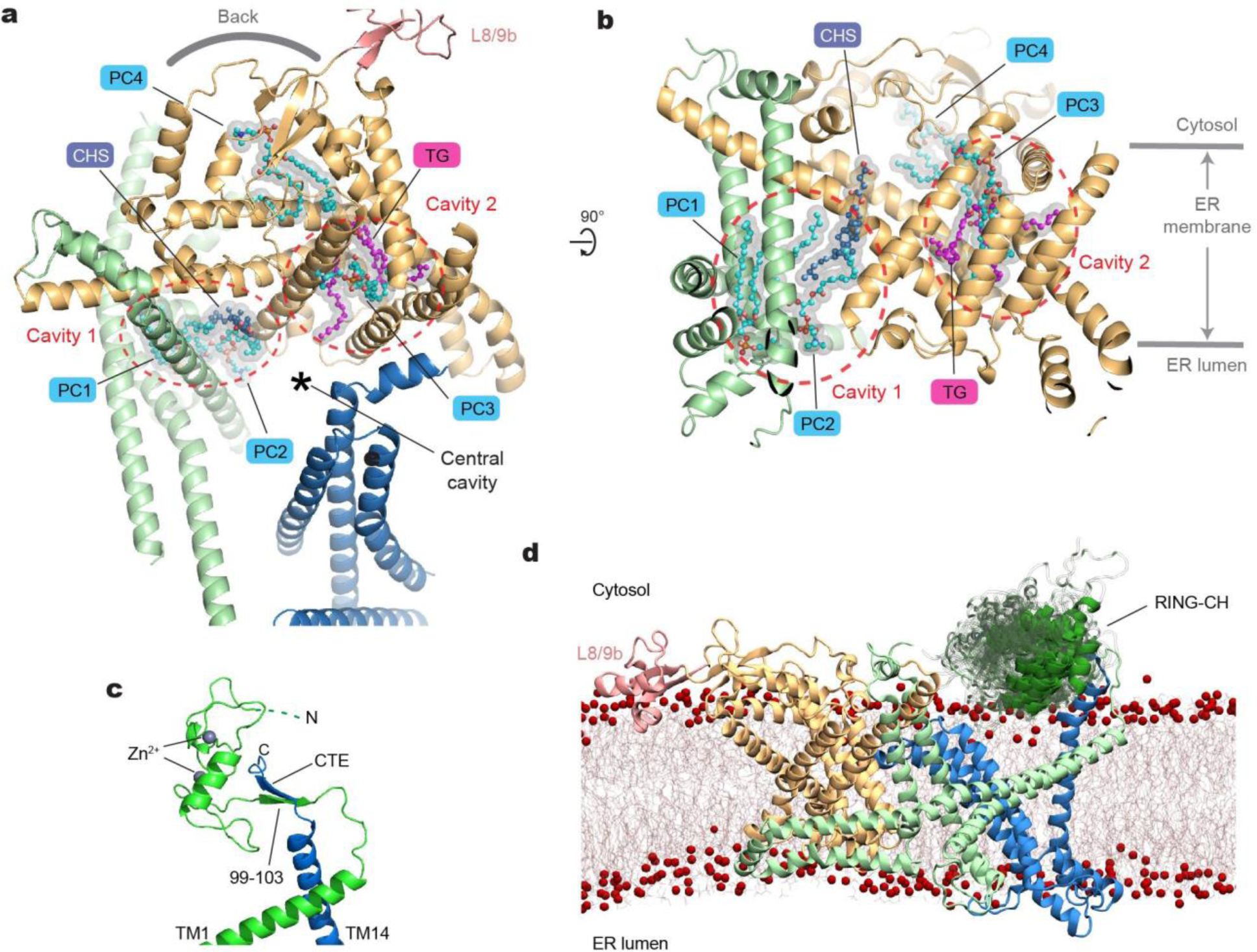
Lipid molecules within the middle domain of Doa10 and the flexibility of the RING-CH domain. **a**, Lipid molecules entrapped in the interior of the TMD. The orientation of the Doa10 model (viewed from the cytosol) is rotated by ∼90° clockwise from Fig. 1b. **b**, As in **a**, but a lateral view from the central cavity to the middle domain. **c**, An AlphaFold2 model of the RING-CH and CTE domains. Shown is a side view, equivalent to view in **d** and Fig. 1f. **d**, A result of an MD simulation on WT Doa10 (side view). Positional flexibility of RING-CH is represented with multiple overlaid structures.

Among the poorly resolved regions in our cryo-EM are the N-terminal RING-CH domain and the C-terminal extension (CTE) following the TM14, which are expected to be in proximity to each other based on our cryo-EM structure (Fig. 1f). AlphaFold2 predicts that the CTE and part of the RING-CH domain co-fold into a two-stranded antiparallel β-sheet (Fig. 2c). Previous biochemical studies found that a truncation of CTE almost completely abolishes the degradation of Deg1 substrates ^35^. This can be explained by a possible loss of proper functioning of the RING-CH domain without the CTE.

To test whether flexibility of the RING-CH domain observed in the cryo-EM structure could be an intrinsic property of Doa10, we ran 1-μs all-atom MD simulations on the hybrid model. Indeed, the results showed markedly larger mobility for the RING-CH domain compared to the TMD (Fig. 2d; Supplementary Fig. 4a–c and Movie 1). Throughout the course of the MD simulations, the CTE remained stably bound to the N-terminal RING-CH domain with the two β-strands forming 4 or 5 hydrogen bonds on average (Supplementary Fig. 4d,e). Thus, this observation suggests that a separation between TMs 1 and 14 is unlikely. Nevertheless, the cryo-EM structure indicates that CTD is relatively more mobile than the rest of the TMD (Supplementary Fig. 3c). Particle classification showed that the CTD wobbles by ∼5° (Supplementary Fig. 4f–h), probably in part due to limited contacts between TMs 1 and 14. This flexibility would allow diffusion of lipid molecules between the bulk membrane and the central cavity of Doa10. On the other hand, integral (even single-spanning) membrane proteins would not easily enter the central cavity laterally through the seam between TMs 1 and 14 as the RING-CH:CTE contact would act as a barrier.

### Structural and sequence conservation of middle domain

Despite the highly interesting architecture of Doa10, the structure itself did not provide clear insights into the functions of the TMD. To better define functionally important features in Doa10, we mapped the amino acid conservation across Doa10 homologs onto the structure (Fig. 3a,b). This shows that the middle domain constitutes the most conserved region, whereas both NTD and CTD are substantially variable. Specifically, within the middle domain, conserved TMs 5, 6, and 7, which together were previously referred to as the ‘TEB4-Doa10 (TD)’ domain, create a ‘tunnel’-like cavity between a wedge formed by the TMs and the roof-like feature formed by the L6/7 loop, together with other parts of the middle domain (Fig. 3b). AlphaFold2 predicts that human homolog MARCH6 also displays a highly similar structure in this region (Supplementary Fig. 5c–e). Among the most conserved residues in Doa10 are those amino acids lining the tunnel.

**Figure 3.**
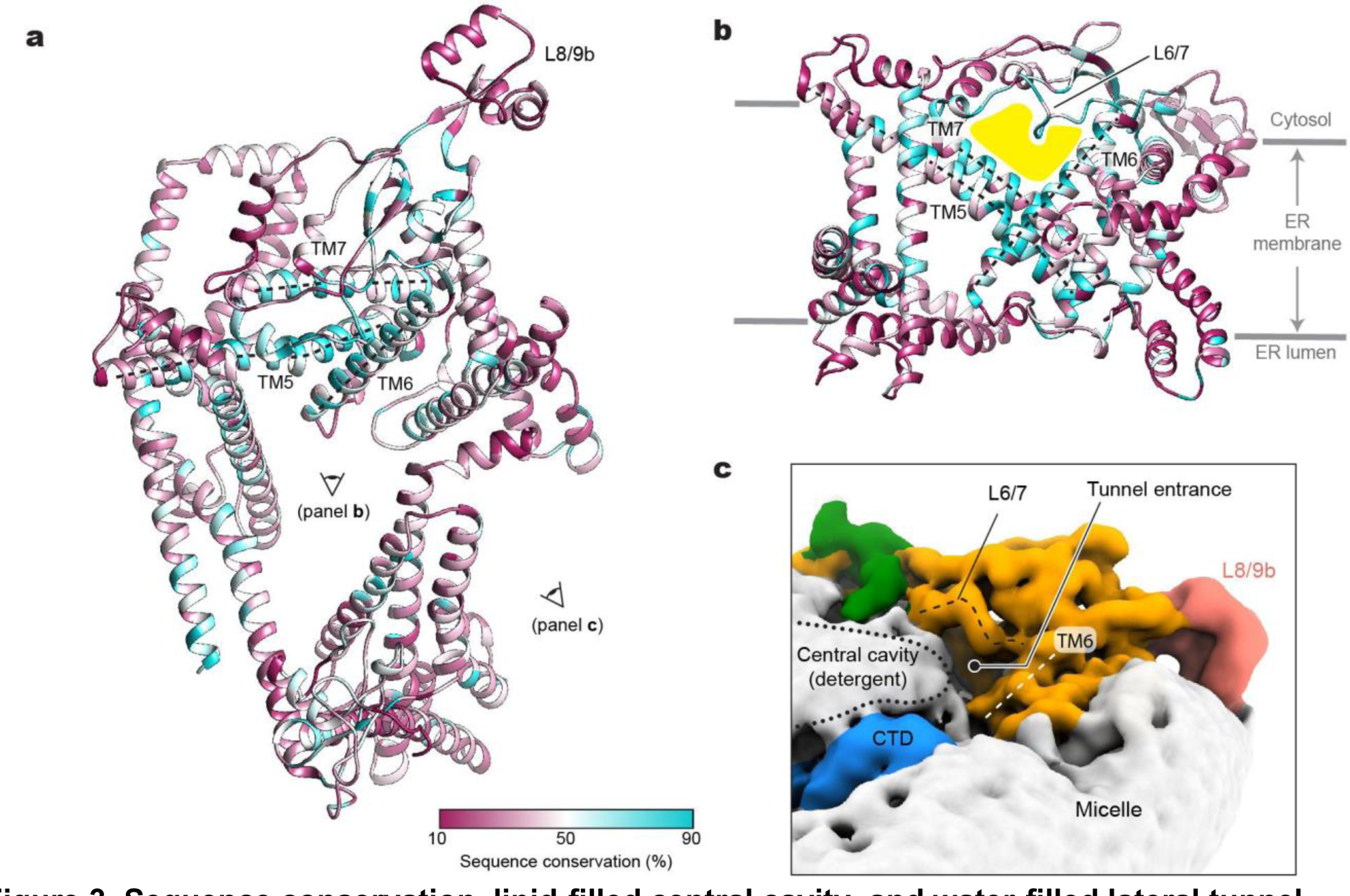
Sequence conservation, lipid-filled central cavity, and water-filled lateral tunnel. **a**, Amino acid sequence conservation is mapped on the structure of Doa10. TMs 5 to 7 (referred to as the TD domain is indicated by dashed lines. The view (cytosolic view) is equivalent to that of Fig. 2a. **b**, As in **a**, but a lateral view from the central cavity to the middle domain. The lateral tunnel is highlighted in yellow. **c**, A view into the entrance of the lateral tunnel. Shown in a lowpass-filtered cryo-EM map. The boundary of detergent features in the central cavity is indicated by a dotted line.

This lateral tunnel, which connects the central cavity and the back of Doa10, is largely level with the cytosolic leaflet of the ER membrane. However, the tunnel interior seems water-filled as the cavity is lined by a mixture of polar, charged, and hydrophobic amino acids. In fact, in our cryo-EM map, the tunnel entrance is exposed to the cytosol with the detergent/lipid density in the central cavity tapering down around the entrance (Fig. 3c). On the other hand, the back of the tunnel seems at least partially blocked by a phospholipid in our cryo-EM structure (PC4 in Fig. 2a,b). Interestingly, one of the acyl chains of this lipid occupies part of the tunnel interior, contributing to hydrophobic surfaces in the tunnel.

Previous biochemical studies have shown that single point mutations of conserved E633 in the tunnel affect the rates of substrate degradation to varying degrees depending on the amino acids it is mutated to (Asp or Gln) and the substrate ^36^. This together with the high sequence conservation and interesting structural features in this region prompted us to further investigate a possible role of the lateral tunnel in substrate recognition.

### Functional importance of the lateral tunnel in the middle domain

To test the effects of mutations in the tunnel on the function of Doa10, we first used the well-established uracil-dependent yeast growth assay ^33^. In this assay, the Ura3 protein is expressed as a Deg1 degron fusion protein in an uracil-auxotrophic (*ura3Δ*) Doa10-null (*doa10Δ*) yeast strain together with an exogenous Doa10 variant. If the expressed Doa10 variant is functional, the growth of yeast in a medium lacking uracil (−Ura) becomes strongly inhibited due to efficient Deg1-Ura3 degradation (Fig. 4a). By contrast, a functionally defective Doa10 variant would allow a growth of the yeast as the Ura3 enzyme remains stable in the cytosol. To increase the dynamic range of the readout, we expressed Doa10 using two promoters with different strengths: *RET2* and *DOA10* promoters. While the *RET2* promoter expresses our exogenous Doa10 constructs at a level comparable to endogenous Doa10 in the WT strain, the expression level from the plasmid-borne *DOA10* promoter was substantially lower than this (Supplementary Fig. 6a), possibly due to the *DOA10* promoter used being partial or because exogenous Doa10 contained many synonymous codon replacements to facilitate molecular cloning of Doa10 (see Methods). We also noticed that a C-terminal GFP-tag somewhat reduces the expression level of Doa10 compared to a shorter peptide tag (ALFA-tag). Using different Doa10 expression levels in this way, we widened the dynamic range of phenotypic readouts of the assay.

**Figure 4.**
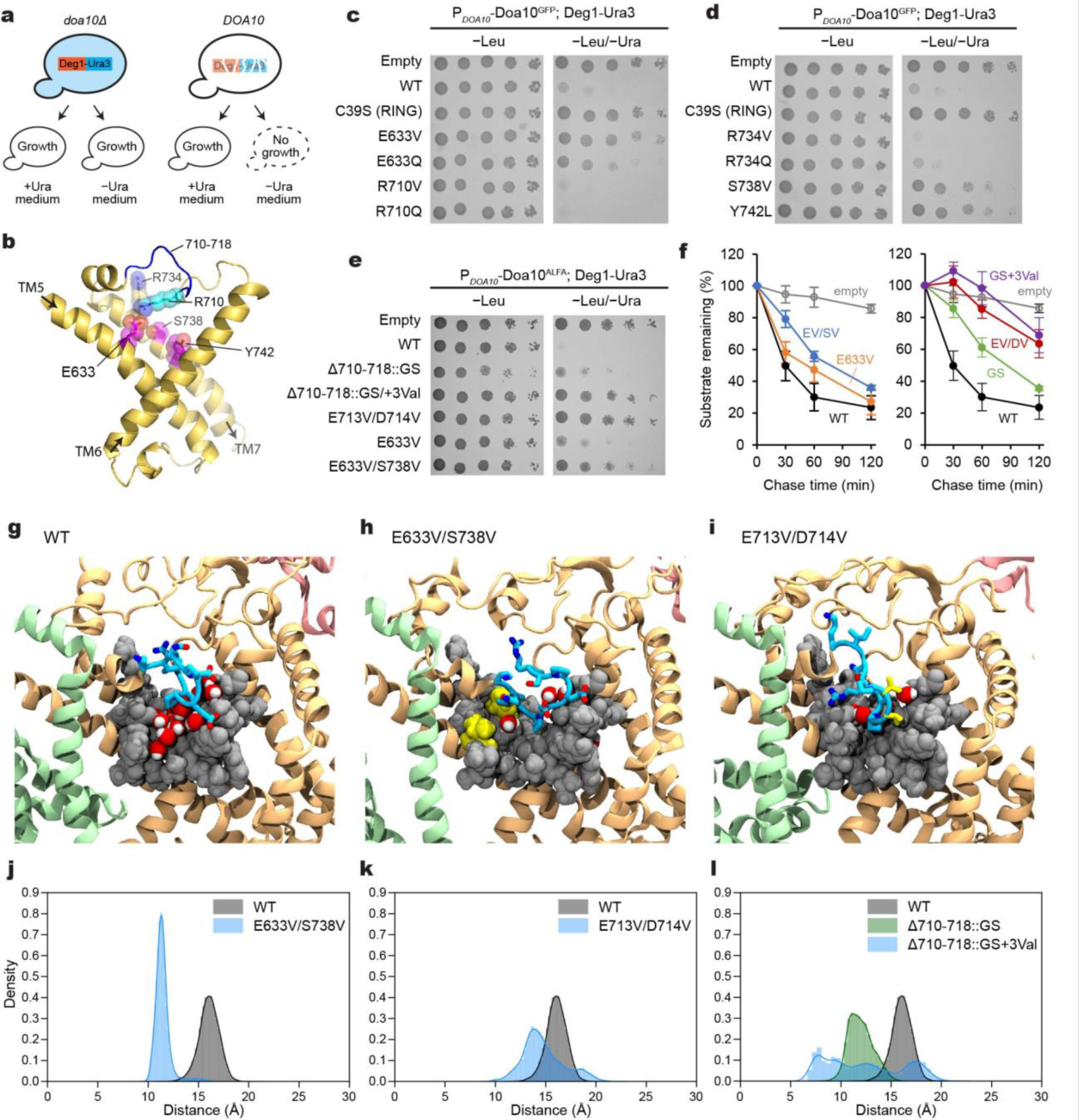
Mutational analysis of the lateral tunnel of Doa10. **a**, Schematic diagram of Doa10-dependent yeast growth inhibition assay using Deg1-Ura3. **b**, Structure of TMs 5 to 7 (TD domain). The view is similar to Fig. 3b. Positions of amino acid residues tested for the effects on Deg1-Ura3 degradation are shown with sticks and spheres (magenta: defects observed; cyan/blue: no substantial defects observed; see **c** and **d**). **c** and **d**, Yeast growth inhibition assay with indicated Doa10 mutants (GFP-tagged Doa10 was expressed at a relatively low level from the DOA10 promoter in a single-copy plasmid). **e**, As in **c** and **d**, but testing different mutants of Doa10 (with an ALFA-tag). Note that a higher expression level of Doa10-ALFA compared to Doa10-GFP (Supplementary Fig. 6a) produces stronger growth inhibition (compare WT and E633V from **c**). **f**, Cycloheximide chase experiments on Deg1-Ura3 using indicated Doa10 mutants (‘EV/SV’=E633V/S738V, ‘GS’=Δ710-718::GS, ‘GS+3Val’=Δ710-718::GS+3Val, and ‘EV/DV’=E713V/D714V). Means and s.e.m. are of three independent experiments. **g**-**i**, Example snapshots of MD simulations on WT and indicated Doa10 mutants. Blue licorice, L6/7 (positions 710-718); gray spheres, space-filling representation of the wedge-shaped tunnel-lining amino acids from TMs 5 to 7; red and white spheres, water molecules within the lateral tunnel; yellow, mutated valine residues. See Supplementary Fig. 7 for GS and GS+3Val mutants. **j**-**l**, The distance distribution was computed between the center of mass of the loop residues (712-715) and the hydrophobic wedge residues (F637, M686, F689, and Y742) for the wild type and mutant systems. The first 150 ns of each trajectory were removed to allow loop relaxation and the 2 replicas were combined. Cyan, residues 710-718; yellow, mutated residues; Gray, van der Waals spheres of wedge-lining residues; red/light gray, water molecules.

When we tested for an E633Q mutant, we observed only a moderate growth rescue as observed previously ^36^ (Fig. 4b–d and Supplementary Fig. 6b), indicating a minor defect in Deg1-Ura3 degradation. The hydrophobic amino acid valine in the same position strengthened this defective phenotype. Mutations on some other, but not all, amino acids lining the tunnel (such as S738V and Y742L) also exhibited similar defects. The double mutant E633V/S738V produced a stronger effect than E633V alone (Fig. 4e), suggesting that increasing the hydrophobicity in the tunnel interior can impair Deg1-Ura3 degradation.

Next, we tested whether altering the roof-like L6/7 feature affects Deg1-Ura3 degradation. A single point mutant on R710 or a larger replacement mutant (replacing Δ710–718 with a glycine/serine linker; Δ710–718::GS) caused no or relatively moderate impairment in Deg-Ura3 degradation. A partial defect phenotype of the latter loop replacement mutant suggests that the native structure of L6/7 is not strictly required for substrate degradation (Fig. 4e and Supplementary Fig. 6c). Strikingly, addition of a few hydrophobic amino acids (Δ710–718::GS+3Val and E713V/D714V) in L6/7 caused a stronger growth rescue. Consistent with this, cycloheximide chase experiments showed substantial stabilization of Deg1-Ura3 with Δ710–718::GS+3Val and E713V/D714V Doa10 (Fig. 4f and Supplementary Fig. 6d–f).

To understand the mechanism underlying the defects caused by the above mutations, we performed 1-μs MD simulations on mutant Doa10 containing E633V/S738V, Δ710–718::GS, Δ710– 718::GS+3Val, or E713V/D714V (Fig. 4g–i; Supplementary Fig. 7 and Movie 2). In all mutants, we observed a collapse of L6/7 onto the wedge-shaped surface of the lateral tunnel formed by TMs 5–7 (Fig. 4j-l). Except for the Δ710–718::GS mutant, this collapse seems to be in part due to increased hydrophobic interactions between the L6/7 and the wedge-shaped surface of the tunnel. Consequently, these mutants exhibited a less solvent-accessible space in the tunnel compared to wild-type Doa10. We note that while the Δ710–718::GS mutation also collapsed L6/7, the interaction pattern was somewhat different from the other mutants tested: the replaced Gly/Ser loop of the Δ710– 718::GS mutant mainly interacted with a polar surface formed by TMs 5 and 7 (Supplementary Fig. 7 and Movie 2), but in other mutants, the loop also interacted with a solvent-exposed hydrophobic surface formed around the bottom tip of the wedge (F637, M686, F689, A690, and Y742) (Fig. 4j-l). If the lateral tunnel is to directly interact with the Deg1 peptide, such a prolonged interaction between the L6/7 and the tunnel’s hydrophobic surface in the mutants would competitively inhibit binding of Deg1, which forms amphipathic helices. On the other hand, the somewhat weaker Deg1-Ura3 degradation defect of the Δ710–718::GS mutant might be explained by the observation that the hydrophobic surface is less stably occupied by its Gly/Ser loop.

### Probing substrate interaction with Doa10 by photocrosslinking

Our structural and mutational analyses suggest the lateral tunnel in the middle domain potentially serves as a substrate-binding site. To probe a direct interaction between Doa10 and the Deg1 degron, we tested UV photocrosslinking between Doa10 and the Deg1 substrate in intact yeast cells. We first incorporated the non-natural photocrosslinkable amino acid p-benzoyl-L-phenylalanine (Bpa) into specific positions within the Deg1 sequence by amber stop codon suppression (Fig. 5a). Three out of four positions in Deg1 yielded crosslink adducts with Doa10 in a UV-dependent manner, suggesting a physical interaction between Deg1 and Doa10 (Fig. 5b).

**Figure 5.**
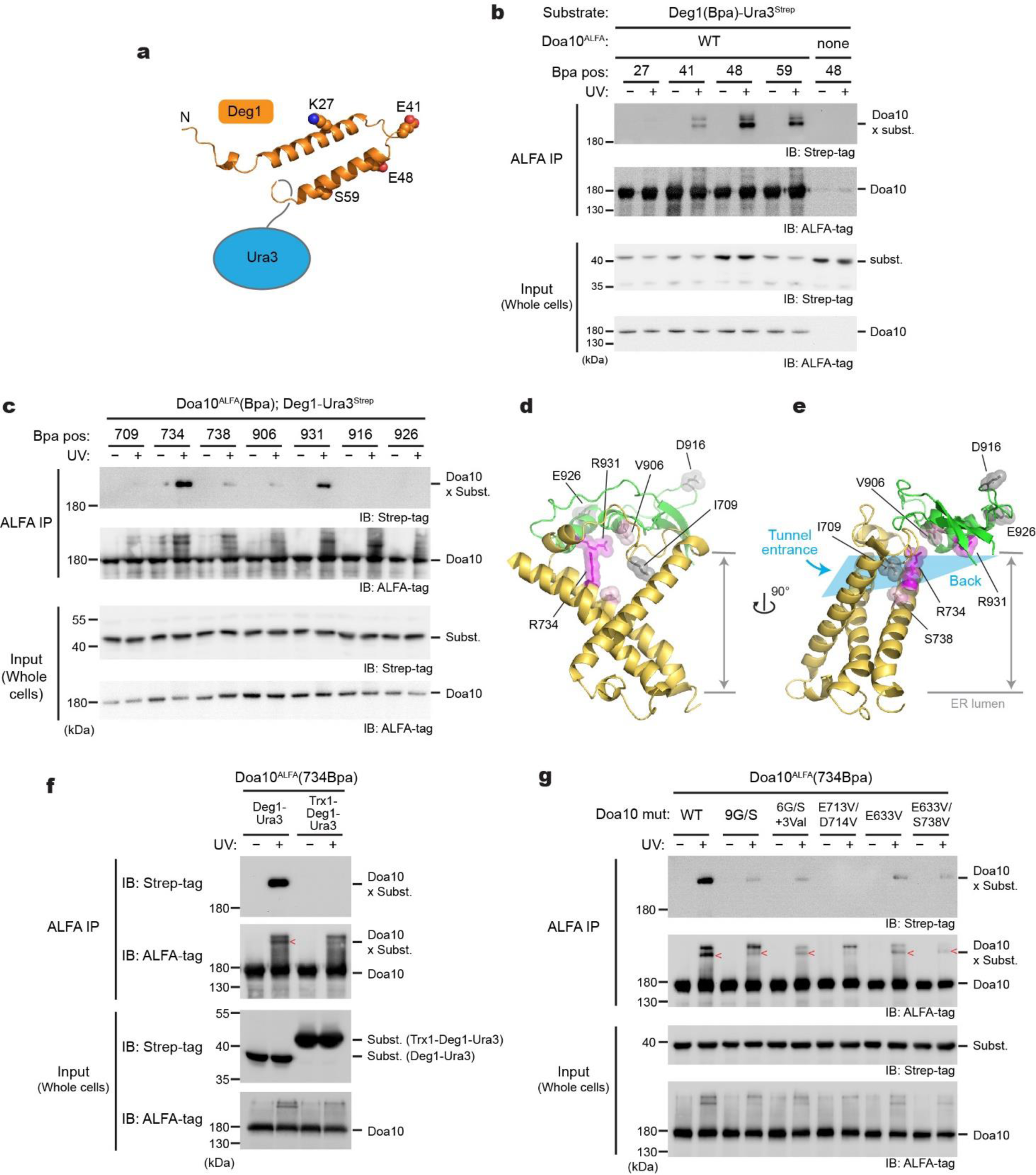
Deg1 inserts into the lateral tunnel of Doa10. **a**, A schematic diagram of Deg1-Ura and the secondary structure of Deg1 (predicted by AlphaFold2). **b**, In-vivo photocrosslinking experiment using Bpa-incorporated Deg1-Ura and wild-type Doa10. IP, immunoprecipitation. **c**, As in **b**, but Bpa was incorporated into a specific position in Doa10 instead of Deg1. **d** and **e**, Structure of the lateral tunnel (**d**, front view; **e**, side view) with highlighting the positions tested for crosslinking in panel **c**. Ribbons in yellow and green are TMs 5–7 and L8/9c, respectively. Residues in magenta crosslink to Deg1-Ura3. **f**, As in **c**, but testing the effects of Trx1 fusion on crosslinking with 734Bpa of Doa10. **g**, As in **c**, but testing the effects of tunnel mutations on crosslinking with 734Bpa of Doa10.

Next, to examine an interaction between the middle-domain tunnel of Doa10 and Deg1, we incorporated Bpa into the tunnel interior and performed crosslinking with Deg1-Ura3 (Fig. 5c). Multiple positions in the tunnel interior showed crosslinking to the substrate (Fig. 5c). Positions 734 and 931 formed strong crosslinking, whereas some other positions (positions 738 and 906) showed weak crosslinking. In contrast, we did not observe crosslinking when we introduced Bpa into the nearby cytosolically exposed surface (positions 916 and 926), demonstrating that the observed crosslinking is not due to random collisions between Doa10 and the substrate. Importantly, the fact that multiple positions in the tunnel interior crosslink with Deg1-Ura3 (Fig. 5d,e) suggests that a part of the substrate polypeptide, most likely the Deg1 peptide, inserts into the lateral tunnel. Considering the confined dimensions of the tunnel, we tested this idea by fusing Trx1, a ∼100-amino-acid-long globular protein, to the N-terminus of the Deg1-Ura3 substrate and performing the crosslinking experiment (Fig. 5f). Indeed, no crosslinking was detected between position 734 of Doa10 and the Trx1-fused substrate, suggesting an inability of Trx1-fused Deg1 to enter the tunnel. Additionally, we examined the effects of mutations in the tunnel and the L6/7 on the Bpa crosslinking. Consistent with the defects observed in growth assays (Fig. 4 e,f), all tested mutations substantially decreased or abolished crosslinking between the tunnel and substrate (Fig. 5g). Collectively, these data show that Doa10’s tunnel interior plays a vital role in facilitating Deg1 recognition through direct interaction.

### Positions of E2

During the substrate polyubiquitination, the RING-CH domain of Doa10 is expected to interact with E2 proteins Ubc6 and Ubc7 to position the C-terminus of the Ub close to the substrate for ligation to occur. Because neither Ubc6 nor the Ubc7–Cue1 complex was included in our cryo-EM analysis, we generated AlphaFold2 models for Doa10 in complex with Ub and Ubc6 or Ubc7–Cue1 (Fig. 6a-b, Supplementary Fig. 8 a and b). These models indeed predicted expected interactions among the RING-CH domain, E2 domain, and Ub. In addition, the models predicted that both Ubc6 and Ubc7– Cue1 associate with Doa10 through interaction with L8/9b of Doa10. Interestingly, Ubc6 and Cue1 are predicted to bind to opposite sides of L8/9b (Fig. 6f), making it possible for both Ubc6 and Ubc7–Cue1 to be simultaneously tethered to Doa10, while the E2 domains of Ubc6 and Ubc7 would engage with the RING-CH domain one at a time.

**Figure 6.**
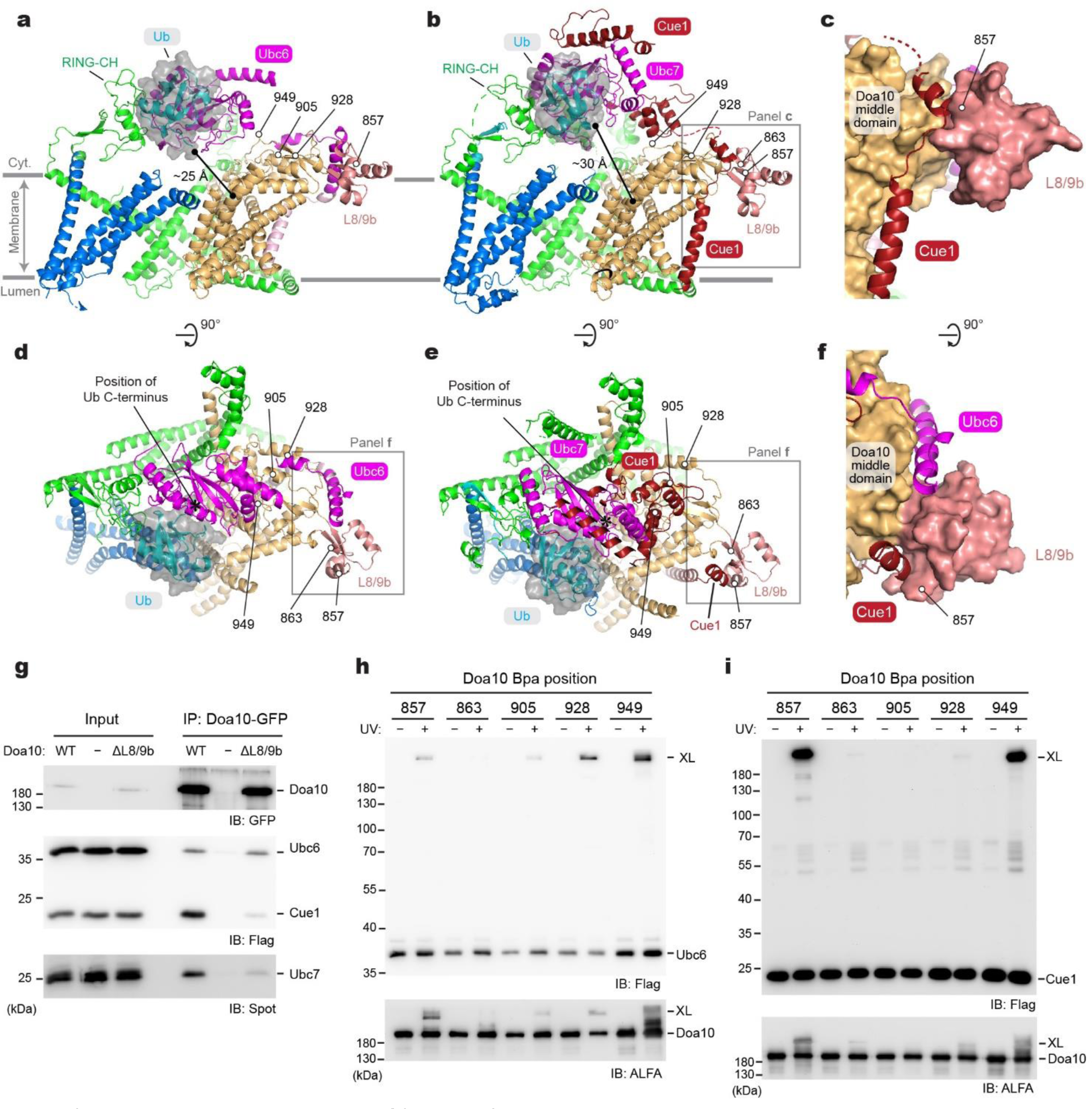
Interaction between Doa10 and E2s and putative polyubiquitination site. **a**, AlphaFold2 model of Doa10 in complex with Ubc6 and Ub (a side view along the membrane plane). For Doa10, the RING-CH and N-terminal domains are shown in green, the middle domain in light brown, L8/9b in salmon, and the C-terminal domain in blue. Ubc6 is shown in magenta, and Ub is represented as cyan ribbon and semitransparent gray surface. **b**, As in a, but showing a model for the Doa10–Cue1–Ubc7–Ub complex. The area outlined with a gray box is shown in panel c. **c**, A view highlighting the interaction between Cue1 and L8/9b of Doa10 (Doa10 is in a surface representation). **d**, As in **a**, but showing the top (cytosolic) view. The area in a gray box is shown in panel f. **e**, As in **b**, but showing the top (cytosolic) view. The area in a gray box is shown in panel **f**. **f**, A view highlighting the interaction of Cue1 or Ubc6 with L8/9b of Doa10 (composite of the two models shown in d and e). **g**, Co-immunoprecipitation of Ubc6, Cue1, and Ubc7 with Doa10-GFP and its L8/9b truncation variant (ΔL8/9b). **h** and **i**, Bpa was introduced to indicated positions in Doa10, and UV photocrosslink to Ubc6 (panel **h**) or Cue1 (panel **i**) was examined by immunoprecipitation and immunoblotting.

To biochemically probe the AlphaFold2-predicted interactions between L8/9b and Ubc6 or Cue1, we first truncated L8/9b from Doa10 (ΔL8/9b) and performed co-immunoprecipitation (co-IP) experiments. Although L8/9b is not a universally conserved feature among the Doa10/MARCH6 proteins (for human MARCH6, see Supplementary Fig. 5), the yeast growth assay indicated that a deletion of L8/9b largely abolishes Deg1 substrate degradation (Supplementary Fig. 8 c and d), as expected for a loss of proper E2 interactions. Consistent with this, the co-IP experiment showed that ΔL8/9b substantially decreased co-purification of Cue1 (Fig. 6g). On the other hand, ΔL8/9b did not affect the amount of co-purified Ubc6, suggesting a possibility that copurified Ubc6 remained bound to Doa10 through an additional interaction site(s).

To further probe Cue1 and Ubc6 interaction with Doa10, we performed in-vivo UV-photocrosslinking experiments by incorporating Bpa into positions in Doa10 based on the AlphaFold2 models (Fig. 6h). Strong crosslink adducts could be observed with multiple positions expected to be close proximity to Ubc6 or Cue1 (positions 928 and 949 for Ubc6, and positions 857 and 949 for Cue1), further providing the evidence for the Doa10-E2 interactions predicted by the AlphaFold2 modeling. However, the observations that position 857 of Doa10 crosslinked to Ubc6 and that no crosslinking was observed between position 863 and Cue1 are somewhat inconsistent with the models.

While further structural investigations would be necessary to fully validate the AlphaFold2 models, an important implication of them is that, in the E2 and Ub-containing complexes, the C-terminal double-Gly tail would be positioned immediately (∼10–15 Å) above the central cavity of Doa10 and in a close (∼25 Å) distance from the tunnel entrance. Thus, the binding of a substrate polypeptide to the lateral tunnel would optimally position the substrate for ubiquitination while flexibility of the RING-CH and E2 domains would allow elongation of a poly-Ub chain. A similar feature of Ub positioning with respect to human MARCH6 could also be found from AlphaFold2 modeling with UBE2J2, a cognate E2 of MARCH6 (Supplementary Fig. 8 f and g)

### Substrate access to the middle-domain tunnel

The well-characterized Deg1 degron provided us with an opportunity to examine requirements in recognition of membrane protein substrates by Doa10. Previously, it has been shown that an insertion of two-spanning membrane protein Vma12 between Deg1 and Ura3 of the Deg1-Ura3 substrate renders much more efficient degradation of the substrate protein ^35^ (Fig. 7a, left panel). The Vma12 portion localizes the protein directly into the ER membrane, and this would increase the frequency of the encounter between Deg1 and Doa10. In our growth assay using Doa10 mutants, we could recapitulate this enhanced degradation of the substrate protein (and thus lesser extents of a growth rescue by mutations) upon Vma12 insertion (Supplementary Fig. 9a).

**Figure 7.**
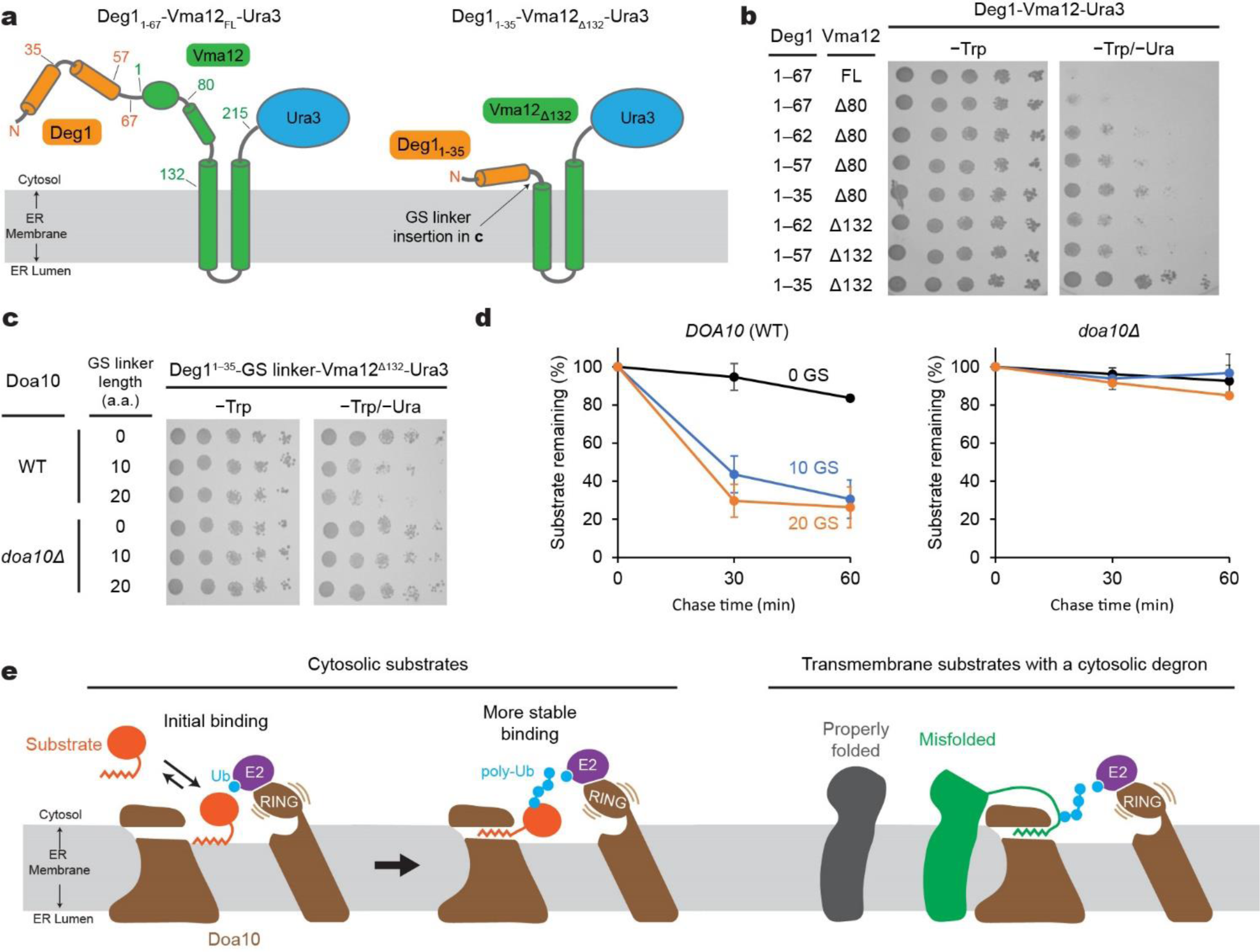
Physical constraints in recognition of a membrane-protein-linked Deg1 degron by Doa10 and working model for substrate recognition and polyubiquitination. **a**, Schematic diagram of the Deg1-Vma12-Ura3 model substrate. Left, original construct; right, minimal construct. Numbers indicate the amino acid positions. **b**, Doa10-dependent yeast growth inhibition using various truncation variants of Deg1-Vma12-Ura3. **c**, As in **b**, but a Gly/Ser (GS) linker was inserted between the truncated Deg1 (Deg1^1–35^) and Vma12^Δ132^. Two different GS linkers were tested: ‘10 GS’ = GGS GGS GGS G and ‘20 GS’= GGS GGS GGS GGS GGS GGS GG. **d**, Degradation of indicated Deg1^1–35^-GS-Vma12^Δ132^ variants were measured by cycloheximide chase and immunoblotting (also see Supplementary Fig. 9b). Means and s.e.m. of three independent experiments. **e**, Working model for the substrate-recognition mechanism of Doa10.

Our model proposing recognition of the Deg1 degron by the lateral tunnel in the middle domain predicts that the Deg1 portion of Deg1-Vma12-Ura3 would also need to access the tunnel interior for efficient polyubiquitination. Because the two TMs of Vma12 must remain in the bulk membrane outside Doa10 while the Deg1 degron interacts with the tunnel, we hypothesized that a certain distance (∼20–30 Å) between Deg1 and the first TM of Vma12 is required for Deg1 recognition. In fact, in the original Deg1-Vma12-Ura3 fusion protein ^35^, there is an ∼130-amino-acid-long segment that can potentially span ∼100 Å between the C-terminus of Deg1 and the first TM of Vma12 (Fig. 7a). To test our hypothesis, we truncated different lengths in the C-terminal region of Deg1 and the N-terminal portion of Vma12 (Fig. 7a,b). In the yeast growth assay, longer truncations in Deg1 and Vma12 resulted in gradually stronger rescue of Doa10-dependent growth inhibition, suggesting that they became less efficient substrates to Doa10. With the complete deletion of the segment of Vma12 preceding the first TM (Δ132) and a minimal Deg1 sequence (residues 1–35), Doa10-dependent degradation of Deg1-Vma12-Ura3 could be almost completely blocked (Fig. 7c,d; Supplementary Fig. 9a). When we reintroduced a flexible Gly/Ser-linker into the truncated region, the degradation was restored both at the steady-state and kinetic levels. Thus, these findings support our model that in the case of membrane-embedded protein substrates, the degron signal needs to be extended out from the membrane domain to reach the lateral tunnel from the periphery of Doa10.

## Discussion

Based on the data presented here, we propose the following model for how Doa10 binds and polyubiquitinates its substrates with a cytosolic degron (Fig. 7e). Our crosslinking experiments showed that Doa10 recognizes degrons, at least Deg1, through a direct interaction within its lateral tunnel formed in the conserved middle domain. This interaction positions the substrates right above the central cavity, which is necessary for efficient polyubiquitination. Many Doa10 degrons characterized so far, including Deg1, contain an amphipathic or hydrophobic motif, and these structural features have been shown to be critical for the Doa10 recognition ^24,28,39–44^. Thus, it is likely that substrate recognition is mediated by hydrophobic interactions between the lateral tunnel and hydrophobic features of the degrons. In addition, prior to insertion into the lateral tunnel, these hydrophobic degrons presumably peripherally associate first with lipids in the central cavity. Such lipid-mediated recruitment of substrates would increase the efficiency of substrate binding to the lateral tunnel as the tunnel is rather concealed near the cytosol-membrane interface. Once the substrate is stably bound, the flexible RING-CH domain and tethered E2s would transfer Ub molecules onto the substrate polypeptide above the central cavity. The horseshoe-like topology of the TMD of Doa10 implies that for targeting membrane-embedded substrates to Doa10, their degrons would need to be extended from their membrane domains. This mechanism might be important for preventing properly folded membrane proteins from non-selectively accessing the lateral tunnel and polyubiquitination site (i.e., central cavity).

Although more substrates of Doa10 remain to be discovered, a current list of substrates suggests that the lateral tunnel can possibly be used for a broad range of substrates besides Deg1. Recently, it has been found that mistargeted TA proteins become substrates of Doa10 after extraction from the mitochondrial outer membrane by Msp1 or from the ER membrane by Spf1 (ref. ^45–47^). Once extracted by these ATPases, these TA transmembrane helices might first peripherally associate with the ER membrane and then be recruited to the central cavity and ultimately to the lateral tunnel of Doa10. It has also been shown that certain unimported mitochondrial matrix proteins are targeted for degradation by Doa10 in addition to two other E3 ligases San1 and Ubr1 in a manner dependent on the presence of an N-terminal mitochondrial-targeting sequence (MTS) ^48^. Given the notion that MTSs typically form an amphipathic α-helix, akin to Deg1, it is tempting to speculate that recognition of these proteins by Doa10 might also be mediated by an interaction between their MTS and the lateral tunnel of Doa10.

Although our data suggest that the lateral tunnel potentially serves as a major recognition site for the ERAD-C-type substrates, it remains to be elucidated whether Doa10 also uses alternative mechanisms for substrate recognition. Beside ERAD-C substrates, Doa10 also can act on certain membrane proteins, such as Sbh2 (a paralog of the Sec61β subunit in yeast) ^28^, which is a TA protein. Doa10-dependent degradation of Sbh2 requires the intact transmembrane helix and the following short ER-luminal C-terminal tail. Thus, Doa10 might first ubiquitinates Sbh2 in a membrane-embedded state, and then Cdc48 might extract polyubiquitinated Sbh2 from the ER membrane. In this scenario, the lateral tunnel, which is oriented toward the cytosol and physically separated from the ER membrane, would be less likely to be used for the recognition of Sbh2.

Another example of ERAD-M-type substrates is Ubc6, which is also a TA protein. Besides functioning as an E2 enzyme for Doa10, it has been shown that Ubc6 can act as a substrate of Doa10 (ref. ^27^). Like Sbh2, it is unclear how Ubc6 is recognized by Doa10 as an ERAD substrate, but it has been shown that a mutation of E633, one of lateral tunnel residues, to aspartic acid greatly reduces the turnover rate of Ubc6 without significantly affecting turnover rates of Deg1 substrates ^36^. A more recent study showed that Doa10 also performs an retrotranslocase activity to extract Ubc6 from the membrane independently of Cdc48 (ref. ^37^). These observations suggest that Doa10 might also use its central cavity and lateral tunnel but potentially in a somewhat different manner for ERAD-M substrates.

In addition to the above-mentioned examples, peripheral membrane proteins, such as the squalene monooxygenase Erg1 and lipid-droplet phospholipase Pgc1, are also on the list of Doa10 substrates ^39,49^. The AlphaFold2 models of Erg1 and Pgc1 suggest that they would peripherally associate with the cytosolic leaflet of the membrane through their C-terminal hydrophobic helices. Since their structures are overall well-folded without an extended tail like Deg1, their degradation is less likely to be dependent on Doa10’s lateral tunnel. Instead, it is possible that Erg1 and Pgc1 may simply enter the area on Doa10’s central cavity from the bulk membrane and become polyubiquitinated. We note that from our cryo-EM structure, the cytosolic leaflet is continuous between the bulk membrane and the central cavity through a space above TMs 1 and 3. Future investigations would be necessary to understand how such peripheral membrane protein substrates are recognized by Doa10 and how the central cavity and lateral tunnel are involved in the process.

Lastly, one highly intriguing feature of Doa10 is the horseshoe-like circular topology, where the N-terminal RING-CH domain co-folds with the CTE at the joint. The topology creates a closed, lipid-filled central cavity. AlphaFold2 models of metazoan MARCH6 and plant SUD1 show that this feature is universal in the Doa10 homologs. Previously, it has been shown that the CTE is critical for Doa10-mediated turnover of most substrates ^35,50^. Why did the RING-CH domain of Doa10 evolve to require partnering with the CTE for its function? One potential function of the CTE might be to restrict the position of the RING-CH domain toward the central cavity. Without additional anchoring through TM14, the RING-CH domain might be too flexible for efficient substrate ubiquitination. Another possibility is to tie the closed circular topology with the RING-CH domain’s activity to prevent other ER integral membrane proteins from accidentally entering the central cavity and being polyubiquitinated. We attempted to create circularly permuted versions of Doa10 that maintain the polyubiquitination function to further test the latter hypothesis, but so far, this effort has been unsuccessful. While our data indirectly suggest that the closed circular topology of Doa10 can act as a barrier for integral membrane proteins, a full understanding of the physiological roles of the circular topology warrants further biochemical studies.

## Methods

### Yeast strains and plasmids

Yeast strains and plasmids used in this study are listed in Supplementary Tables 2 and 3.

To enable purification of Doa10 from yeast, we modified the yeast strain BY4741 by inserting a sequence encoding a cleavable GFP-tag between the last amino acid and stop codons of the chromosomal copy of Doa10. The final expressed protein from the resulting strain ySI-118 has an amino acid sequence of (N-terminus)…ENLPDES (Doa10)-AGGATTASGTG (linker)-ENLYFQG (a tobacco etch virus [TEV] protease site)-TASGGGS (linker) KGEELF…(GFP)-(C-terminus). A sequence containing the TEV-GFP-tag and a nourseothricin resistance marker module was amplified from pSK-B399-GFP-NAT (a gift from the Klinge lab) with primers containing homologous arms to the chromosomal insertion site (forward primer: acgaggtttacactaagggtagagctttagaaaatttaccagatgaaagtGCTGGAGGGGCTACCACG; reverse primer: aacatataacttaatgtagatatatatatgtaaatatgctagcattcattGGCCGCATAGGCCACTAG. Lowercase for a homologous sequence to the Doa10 sequence, and uppercase for binding to the plasmid pSK-B399-GFP-NAT). The amplicon was transformed to the BY4741 strain using standard lithium acetate protocol and plated on a YPD agar plate (1% yeast extract, 2% peptone, 2% glucose, and 2% bacto-agar) supplemented with 100 µg ml^-1^ nourseothricin. Single colonies were grown with a YPD medium under nourseothricin selection, and correct insertion was verified using PCR of genomic DNA and by Sanger sequencing.

To overexpress the GFP-tagged Doa10, the endogenous promoter was replaced with a *LEU2*::P*_GAL1_* cassette that was assembled using overlap extension PCR. The *LEU2* expression cassette was amplified from pYTK075 (forward primer: gctaagataatggggTCGAGGAGAACTTCTAGTATATCTAC, lowercase for a homologous sequence to GAL1, uppercase for binding to 5’ end of the LEU1 gene. Reverse Primer: gatttcaaaaactgtttttttagccaagagtaccactaattgaatcaaagCTGCCTATTTAACGCCAAC, lowercase for a homologous sequence to the chromosomal sequence flanking Doa10’s promoter and uppercase for binding to LEU2 gene). P*_GAL1_* was amplified from pYTK030 (Forward primer agaagttctcctcgaCCCCATTATCTTAGCCTAAAAAAAC, lowercase for a homologous sequence to LEU2, uppercase binds to 5’ end of the *GAL1* gene. Reverse primer: tgcagttcatctcttaacctggagacattaacgtcagaatcaacatccatTATAGTTTTTTCTCCTTGACGTTAAAGTATAG, lowercase for a homologous sequence to the chromosomal sequence flanking Doa10’s promoter and uppercase for binding to *GAL1* gene). pYTK plasmids are part of MoClo Yeast Tool Kit ^51^. The two PCR products were purified and amplified as overlapping PCR with the terminal primers. The final amplicon was transformed into yeast strain ySI-118, and selected on SC(−Leu) agar medium containing 2% glucose. Single colonies were selected, and proper integration was verified using PCR of genomic DNA. The resulting strain is designated ySI-154.

To generate Doa10-expressing plasmids for mutational and crosslinking studies, we adapted a Golden Gate cloning strategy ^52^. Full-length Doa10 nucleotide sequence is known to be toxic to *E. coli* cells, preventing propagation of plasmids containing the coding sequence (CDS) of Doa10 (ref. ^53^). To overcome this issue, we splitted the CDS of Doa10 into two plasmids, which did not exhibit growth inhibition in *E. coli* cells. The first plasmid contains a promoter for Doa10 expression (either the endogenous *DOA10* promoter, a *GAL1* promoter, a *RET2* promoter or a *TDH3* promoter), the first 1824 bp of the CDS of Doa10, two BsaI endonuclease restriction sites, and a CEN6-ARS4 module, in this order. The second plasmid contains a BsaI site, 1822th to 3957th bp of the CDS of Doa10 and a C-terminal tag (either GFP or ALFA-tag), an *ENO1* terminator sequence, and *LEU2* marker cassette, and a second BsaI site. The full-length Doa10-encoding plasmid was formed by joining the two plasmids using BsaI Golden Gate cloning prior to transformation into yeast cells. Transformation of yeast with each split plasmid alone does not form colonies on a Leu drop-out agar medium because of separation of the *LEU2* marker and CEN6-ARS4. Only the full plasmid joined by a BsaI Golden Gate reaction, which encodes a full length Doa10 CDS, can be stably maintained in yeast cells. Proper expression of Doa10 proteins were confirmed by immunoblotting analysis of whole cell lysates of transformants. We note that the DNA sequence for the first 322 amino acid residues of Doa10 in the first plasmid originated from gene synthesis of a reverse translated sequence and thus contains many silent mutations.

For the yeast growth (spot) assays using Deg1-Ura3 as a substrate, yeast strain MHY4086 (*doa10Δ*::hphMX4, *lys2-801*::*LYS2*::Deg1-Ura3) was used ^54^. Plasmids with various Doa10 mutants were generated by the Golden Gate cloning approach as described above and transformed into MHY4086. Doa10 was expressed either from an endogenous promoter (P*_DOA10_*) or *RET2* promoter (P*_RET2_*). Colonies were selected on a −Leu drop-out synthetic complete medium (SC[−Leu]) containing 2% glucose.

For the spot assays using Deg1-Vma12-Ura3 as a substrate, yeast strain MHY10818 (*doa10Δ*::*hphMX*; ref. ^50^). The strain was first transformed with plasmid p414-Deg1-Vma12-Ura3 (ref. ^33^), which constitutively expresses Deg1-Vma12-Ura3 under a *MET25* promoter and uses a Trp auxotroph marker for selection. The resulting strain was transformed with a plasmid expressing a Doa10 variant as described above. Colonies were selected on SC(−Trp/−Leu) agar medium containing 2% glucose. All the truncated and GS linker versions of Deg1-Vma12-Ura3 were generated by PCR using the plasmid p414-Deg1-Vma12-Ura3 as a template. The original plasmid p414-Deg1-Vma12-Ura3 contains a FLAG-tag between Deg1 and Vma12, which was removed in the truncated versions. To detect the substrate, we attached a FLAG-tag to the C-terminus of Ura3. All these plasmids were then transformed into either JY103 (wild-type *DOA10*) or MHY10818 (*doa10Δ*).

For the cycloheximide chase experiment using Deg1-Ura3 as a substrate, the yeast strain yKW-283 (*doa10Δ*::*natMX*; *leu2*::P_PGK1_-Deg1-Ura3-2xstrep::*hphMX4)* was used. This strain was made from ySI-167 (*doa10Δ::natMX*) by integrating P*_PGK1_*-Deg1-Ura3-Strep into the *leu2* locus. Strain ySI-167 was made by deleting chromosomal *DOA10* gene (*doa10Δ*::*natMX*) in BY4741 via transformation of a PCR product, which was amplified from pSK-B399-GFP-natMX (forward primer: gatttcaaaaactgtttttttagccaagagtaccactaattgaatcaaagCTGTTTAGCTTGCCTCGTCC; Reverse primer: aacatataacttaatgtagatatatatatgtaaatatgctagcattcattGGCCGCATAGGCCACTAG. Lowercase for a homologous sequence to the *DOA10* gene and uppercase for binding to the natMX marker; pSK-B399 is a gift from the Klinge lab). The amplicon was transformed into BY4741, and selected on YPD agar plate supplemented with 100 µg ml^-1^ nourseothricin. Chromosomal deletion was verified using PCR of genomic DNA. To generate yKW-283, we first generated an integration plasmid (pKW155), expressing Deg1-Ura3-2xstrep under the *PGK1* promoter, using the MoClo Yeast Tool Kit (YTK). The sequence of Deg1 and Ura3 was amplified and cloned individually into a pYTK001 entry plasmid, resulting pKW043 (pYTK001-Deg1), pKW050 (pYTK001-Ura3). Subsequently, all the part plasmids, including pYTK-011 (P*_PGK1_*), pKW043 (Deg1), pKW050 (Ura3 CDS), pYTK-e205 (twin Strep-tag) and pYTK-061 (*ENO1* transcription terminator), were assembled into pYTK-e102 (an integration plasmid targeting to the *LEU2* locus with a hygromycin marker) by BsaI Golden Gate assembly, resulting pKW155. pKW155 was linearized with *Not*I and was transformed into ySI-167. Colonies were selected on YPD agar medium supplemented with 400 µg ml^-1^ hygromycin B. Plasmids with various Doa10 mutants were generated by *Bsa*I Golden Gate cloning approach as described above, and transformed into yKW-283. Colonies were selected on SC(−Leu) agar medium containing 2% glucose.

For the site-specific photo-crosslinking experiments, yeast strain ySI-266 (*doa10Δ::hphMX4 cue1Δ*::*natMX*) was first made by deleting chromosomal *CUE1* (*cue1Δ::natMX*) in MHY10818 via transformation of a PCR product, which was amplified from pYTK078 (*natMX* marker) (forward primer: cgccataaagcattacaatctacgatcgcgcaaacttttttcttttggccCTGTTTAGCTTGCCTCGTCC; Reverse primer: ttatgcgcattatgggcacacttgcgtgttcccggtaagcacttaagcgtGGCCGCATAGGCCACTAG. Lowercase for a homologous sequence to the Cue1 gene and uppercase for binding to the natMX marker). Chromosomal deletion was confirmed using PCR of genomic DNA. Subsequently, ySI-266 was transformed with SNRtRNA-pBpaRS(*TRP*) (ref. ^55^). Colonies were selected on SC(−Trp) agar medium containing 2% glucose. The resulting strain was then transformed with a plasmid expressing Deg1-Ura3, which was generated by assembling pYTK-009 (*TDH3* promoter), pKW043 (Deg1), pKW050 (Ura3), pYTK-205 (twin Strep-tag) and pYTK061 (*ENO1* terminator) into pYTK-e11 (a CEN/ARS plasmid with a Ura3 marker). Colonies were then selected on SC(−Trp/−Ura) agar medium containing 2% glucose. Subsequently, plasmids expressing P*_TDH3_*-Doa10-ALFA with an amber codon mutation at various sites were generated by site-directed mutagenesis and then *BsaI* Golden Gate cloning. Colonies were selected on SC(−Trp/−Ura/−Leu) agar medium containing 2% glucose.

For co-immunoprecipitation of E2 enzymes with Doa10, a plasmid (pYC-302) expressing Cue1, Ubc7 and Ubc6 was made. The chromosomal *CUE1*, *UBC6* and *UBC7* were first amplified and cloned into pYTK-001 entry plasmids individually. The resulting plasmids were then assembled using YTK parts to add a *GAL1* promoter, an epitope-tag, and an *ENO1* terminator by Golden Gate cloning and to form the multigene-expression plasmid pYC-302 (P*_GAL1_*-Cue1-2xFLAG | P*_GAL1_*-Ubc7-2xSPOT | P*_GAL1_*-3xFLAG-6xHis-Ubc6; a CEN/ARS plasmid with a Ura3 selection marker).

For photo-crosslinking experiments with Cue1 and Ubc6, plasmids pYC-300 and pYC-301 were generated to express Cue1 and Ubc7 and Ubc6 respectively. The pYTK-001 plasmids coding *CUE1*, *UBC6* and *UBC7* were assembled by Golden Gate cloning to form pYC-300 (P*_GAL1_*-Cue1-2xFLAG | P*_GAL1_*-Ubc7-2xStrep; a CEN/ARS plasmid with a Ura3 selection marker) and pYC-301 (e111-P*_GAL1_*-3xFLAG-6xHis-Ubc6; a CEN/ARS plasmid with a Ura3 selection marker) as described above. Yeast strain ySI-266 (*doa10Δ::hphMX4 cue1Δ*::*natMX*) was used for Cue1 crosslinking, while yYC-307 (*doa10Δ::hphMX4 ubc6Δ*::*natMX*) was made using a similar strategy to delete chromosomal *UBC6* (*ubc6Δ::natMX*) (forward primer: gactttaaatattaactaaaaccgcattcgcaaattgcaaacaaagtacgtacaatagtaCTGTGGATAACCGTAGTCG; Reverse primer: tcaaaatttatctaaagtttagttcatttaatggcttcatttcataaaaaggccaaccaaGGGCGTTTTTTATTGGTC. Lowercase for a homologous sequence to the *CUE1* gene and uppercase for binding to the natMX marker sequence in the YTK plasmid pYTK078.)

### Purification of Doa10 protein

Yeast strain ySI-154 was inoculated into a YP-raffinose medium (1% yeast extract, 2% peptone, and 2% raffinose) to an optical density at 600 nm (OD_600_) of 0.2 and grown in shaker flasks at 30°C until OD_600_ reached 0.5. Doa10 expression was then induced by adding 2% galactose to the culture, and cells were grown until OD_600_ reached ∼2. Cells were then pelleted, flash frozen in liquid nitrogen, and stored in −75°C until purification.

Cell lysis was performed by cryo-milling (SPEX SamplePrep) cycling 15 times with 1 min on time and 2 min off time. Broken cells were resuspended in a buffer containing 50 mM Tris-HCl pH 7.5, 200 mM NaCl, 1 mM EDTA, 10% glycerol, 2 mM DTT, supplemented with protease inhibitors (5 µg/ml aprotinin, 5 µg/ml leupeptin, 1 µg/ml pepstatin A, and 1.2 mM PMSF). Membranes were further solubilized by the addition of 1% lauryl maltose neopentyl glycol (LMNG; Anatrace) and 0.2% cholesteryl hemisuccinate (CHS; Anatrace), and stirring for 1.5 h at 4°C. The cell lysate was clarified by ultracentrifugation using Beckman Type 45 Ti rotor at 40,000 RPM for 1 hr. The clarified lysate was supplemented with 25 μg/mL Benzonase nuclease and incubated with home-made agarose beads conjugated with anti-GFP nanobody at 4°C for 2.5 hr by gentle rotating. The sample was transferred to a gravity column, washed with a buffer (WB) containing 50 mM Tris-HCl pH 7.5, 200mM NaCl, 1.0 mM EDTA, 2 mM DTT, 0.02% glycol-diosgenin (GDN; Anatrace), and 10% glycerol. Bound Doa10 was eluted by incubating the beads with ∼10 μg/mL TEV protease overnight. The eluate was then injected to a Superose 6 10/300 GL Increase column (GE Lifesciences) equilibrated with 20 mM Tris-HCl pH 7.5, 100 mM NaCl, 1mM EDTA, 2 mM DTT, and 0.02% GDN. Peak fractions were concentrated to ∼5.5 mg/mL using Amicon Ultra (100-kDa cutoff; GE Lifesciences) and used immediately for cryo-EM grid preparation.

### Cryo-EM analysis

The Doa10 sample was supplemented with 3 mM FFC8 (Anatrace) before freezing cryo-EM grids. 3 μL of the sample was applied on each gold Quantifoil R 1.2/1.3 holey carbon grid that was glow discharged for 35 s using a PELCO easiGlow glow discharge cleaner. The grids were blotted for 3-4 s using Whatman No. 1 filter paper at 4 °C and 95-100% relative humidity, and plunge-frozen into liquid ethane using a Vitrobot Mark IV (FEI company). Data was collected on a Krios G2 microscope (FEI company) equipped with a Gatan Quantum Image Filter (with 20 eV slit width) and a Gatan K3 direct electron detector (Gatan). The microscope was operated at an acceleration voltage of 300 kV. The magnification was set to 64,000× under the super-resolution mode with a physical pixel size of 0.91 Å. The total exposure was set to 50 electrons/Å^2^ divided into 50 frames, and the defocus range was set between −0.8 and −2.0 µm. All the data was acquired using SerialEM software.

For detailed illustration of the data analysis, refer to Supplementary Fig. 2c. In short, two datasets, 1,760 movies (Dataset 1) and 2,679 movies (Dataset 2), were pre-processed first with Warp (version 1.0.9; ref. ^56^) to produce an initial model that was used for cryoSPARC template picking, and again in CryoSPARC (version 2.15.0; ref. ^57^) to produce the final set of particles for 3D reconstruction. In Warp, the movies were corrected for motion, and contrast transfer function (CTF) estimated on micrographs divided into 7 × 5 tiles, and particles were picked by Warp’s BoxNet algorithm yielding 299,741 and 241,653 autopicked particles for Dataset 1 and Dataset 2, respectively. The particles in each dataset were imported to CryoSPARC for 2D classification and ab-initio reconstruction, generating three initial models. Only one of the 3 maps presented proteinaceous features and was selected as a template for CryoSPARC template particle picking. The raw movies were re-processed using CryoSPARC (tile-based motion correction, CTF estimation, and manual movie curation) and a new particle data set containing 455,702 and 613,378 particles was obtained with the template picker. The datasets were subjected to a 2D classification, and good classes were selected after visual inspection. For Dataset 1, we used the previously obtained ab-initio models to generate three heterogeneous refinement models, which were subsequently used as reference for the heterogeneous refinement of Dataset 2. A single 3D class from each data set was selected and their corresponding particles were subjected to a second round of heterogeneous refinement with the same 3D reference models. The particles corresponding to the best class in each data set were combined, and after a CTF refinement, particle curation yielded 324,019 particles. A non-uniform refinement job (CryoSPARC version 3.0.0) on these particles resulted in the final map of Doa10 at 3.2-Å overall resolution. The enhanced map shown in Fig. 1 d–e was generated using DeepEMhancer (ref. ^58^). The local resolution distribution is calculated in CryoSPARC.

### Atomic model building

An initial atomic model was generated de novo using Coot (version 0.9) and cryo-EM maps that were sharpened by various B-factors. The model was then refined using the real-space refinement program of the Phenix package (version 1.16; ref) and a cryo-EM map sharpened at a sharpening B-factor of −85 Å^−2^. During the course of this study, AlphaFold2 was published ^38^. The AlphaFold2 model (https://alphafold.ebi.ac.uk/entry/P40318) enabled us to build additional amino acids in CTD (TMs 11-14) and L8/9b, which were difficult to confidently register de novo due to their lower local resolution in our cryo-EM map. The final model was refined again with real-space refinement of Phenix (version 1.19.2). Molprobity in the Phenix package was used for the validation. Structural models for Doa10 with E2s and Ub were predicted with AlphaFold2 (version 2.1.1) using full-length amino acid sequences of the proteins. Figures of cryo-EM maps and atomic models were generated using UCSF Chimera (ref. ^59^), ChimeraX (ref. ^60^), and PyMOL (Schrödinger)

### Sequence conservation analysis

Amino acid sequences of Doa10 from various species were obtained from UniRef90 (https://www.uniprot.org). Out of 100 sequences available, 14 sequences that were shorter than 800 amino acids were removed, and the remaining 86 sequences were subjected to multiple sequence alignment using MAFFT (https://www.ebi.ac.uk/Tools/msa/mafft/) with default parameters. Aligned sequences were opened in UCSF Chimera to map amino acid conservation onto the Doa10 structure.

### Molecular dynamics (MD) simulations

The Doa10 MD simulations were based on the hybrid atomic model described above. In addition to protein, densities were observed that can be fit with 4 phosphatidylcholine molecules, a triglyceride, an ergosterol, and a cholesterol-like detergent. These were modeled with 4 POPC molecules, a triglyceride (16:0,16:1,18:0), an ergosterol, and the detergent was replaced with ergosterol. Zinc ions were added to the RING-CH domain, with coordinating cysteine residues deprotonated and no further constraints applied ^61^. Otherwise, residue protonation states were assigned with H++ (ref. ^62^). The N terminus, starting at residue 30, and C terminus, residue 1318, were capped with neutral acetyl and amide groups, respectively. The termini for the missing loop residues were also neutral capped. This WT model was placed in a realistic yeast ER membrane consisting of 48% POPC, 20% POPE, 10% PLPI, 8% POPS, 3% POPA, 10% ERG, and 1% DYGL (ref. ^63–65^) using CHARMM-GUI (version 3.7 accessed May 2023; ref. ^66^). Double mutants (E633V/S738V, E713V/D714V) were generated by mutating the sidechains of the respective residues. Loop substitution mutants were generated by replacing 710-718 with the following sequences: GGSGGSGGS (Δ710-718::GS) or GGSVVVGGS (Δ710-718::GS+3Val). The wild type and mutant systems were similarly placed in the yeast ER membrane model, hydrated using a TIP3P (ref. ^67^) water box, and neutralized with 0.15 M KCl. The all-atom systems were ∼408,000 atoms. The CHARMM36m protein ^68^ and CHARMM36 (ref. ^69^) force fields were employed in all simulations.

Each system was equilibrated in stages using the following protocol; 1) an initial minimization was performed followed by 2) relaxation of the lipid acyl chains for 1 ns with position restraints applied to all other atoms, 3) 10 ns with the protein and bound lipids restrained to their starting positions, and 4) 100ns with only the protein backbone restrained to allow for lipid relaxation about the protein. Finally, an additional minimization (2000 steps) prior to unrestrained NPT dynamics was performed. For equilibration, NAMD 2.14 was used, while production runs were performed in duplicate for 1 μs per replica with GPU-accelerated NAMD3 (ref. ^70^). All simulations were performed at a constant temperature of 310 K using Langevin dynamics (damping coefficient 1/ps), a constant pressure of 1 atm using a Langevin piston barostat, and periodic boundary conditions. Following initial equilibration, hydrogen mass repartitioning was invoked, allowing for a 4-fs time step ^71^. Short range non-bonded interactions were cut off at 12 Å, with a force-based switching function starting at 10 Å. Long range non-bonded interactions were calculated using particle-mesh Ewald method with grid spacing of at least 1/Å^3^ (ref.^72^). Analysis was carried out and images were rendered with VMD (version 1.9.4a51; ref. ^73^).

### Yeast growth assay

Overnight cultures were diluted in a five-fold serial dilution from an OD_600_ of 0.1, and 5 µl each were spotted onto an indicated agar medium. Plates were incubated at 30°C for 2-3 days before imaging.

### Cycloheximide chase assay

For the experiment in Fig. 4f, the yeast strain yKW-283 (*doa10Δ leu2*::P*_PGK1_*-Deg1-Ura3-strep) was transformed with an empty vector (pYTK-e112) or plasmids encoding various Doa10 mutants under a *DOA10* promoter. Cells were grown in a synthetic medium SC(−Leu). For the experiment in Fig. 7d, the yeast strains were identical with the ones used in the yeast growth assays in Fig. 7c and grown in a synthetic medium SC(−Trp).

Overnight cultures were diluted to an OD_600_ of 0.2 and grown at 30°C until the cells reached mid-log growth phase. 5 OD_600_ units of cells were collected. The pellets were resuspended in 4 ml of fresh medium. 0.25 mg/ml cycloheximide was added to the yeast suspension. Transferred 950 µl of the yeast suspension to a tube containing 20x stop mix (200 mM sodium azide and 5 mg/ml BSA). Pelleted the cells by centrifugation at 6000 *g* for 1 min. The pellets were resuspended with 200 µl of 0.1 M NaOH and incubated at room temperature for 5 min. Subsequently, cells were spun down by centrifugation at 12000 *g* for 1 min and resuspended in 50 ml of reduced SDS sample buffer. Samples were heated at 95°C for 5 min before analysis by SDS-PAGE and immunoblotting.

### Site-specific photo-crosslinking assay

In all the substrate crosslinking experiments, the yeast strain ySI-266 was transformed with the plasmid SNRtRNA-pBpaRS(*TRP*), the plasmid expressing Deg1-Ura3-Strep under the *TDH3* promoter, and the plasmid expressing ALFA-tagged Doa10 under the *TDH3* promoter. An amber codon was introduced at a specific site using site-directed mutagenesis.

Overnight culture was diluted to an OD_600_=0.02 in 50 ml of fresh minimal medium SC(−Trp/−Leu/−Ura) containing 2% glucose and 2 mM Bpa (Amatek, cat# A-0067). The cells were grown at 30°C overnight until the cells reached an OD_600_ of 1.0 to 1.2. Cells were harvested, washed with deionized water and aliquoted into two tubes. One was kept on ice as a control experiment, while the other sample was transferred to a 24-well plate and UV irradiated for 1 h on ice-cold water in the cold room. Cells were pelleted and resuspended in 0.5 ml of lysis buffer (buffer LB, 50 mM Tris-HCl pH7.5, 200 mM NaCl, 10% glycerol, 2mM DTT, 1mM EDTA) supplemented protease inhibitors (5 µg ml^−1^ aprotinin, 5 µg ml^−1^ leupeptin, 1 µg ml^−1^ pepstatin A, and 1 mM PMSF). Cells were then lysed by beating with 0.5-mm glass beads. Cell lysate was supplemented with 1% LMNG and 0.2% CHS and incubated at 4°C for 1 h to solubilize membranes. Subsequently, the lysate was clarified by centrifugation at 17,000 *g* for 30 min. The supernatant was incubated with Sepharose beads conjugated with anti-ALFA nanobody at 4°C for 1 h. The beads were washed three times with 1 ml of wash buffer (buffer WB, 100mM NaCl, 20mM Tris-HCl pH 7.5, 1mM EDTA, 2 mM DTT and 0.02% DDM/CHS). Samples were eluted by addition of reduced SDS sample buffer and mildly heated at 37°C for 30 min before analysis by SDS-PAGE and immunoblotting.

For crosslinking between Doa10 and E2 enzymes, yeast strain ySI-266 (for Cue1) or yYC-307 (for Ubc6) was subsequently transformed with SNRtRNA-pBpaRS, the E2-expression plasmid (pYC-300 for Cue1, pYC-301 for Ubc6), and the plasmid encoding a Doa10 amber codon mutant under the *TDH3* promoter. The cells were grown and treated with the same procedure as described above.

For urea wash controls of photocrosslinking adducts, after washing beads once with buffer WB, two additional washes were performed with buffer WB containing 6 M urea and 0.5% Triton X-100 (buffer WD). This was followed by another wash with buffer WB without urea and elution with SDS sample buffer.

### Co-immunoprecipitation

For co-immunoprecipitation of E2 with Doa10 (Fig. 6g), yeast strain ySI-167 was first transformed with pYC-302, and subsequently with an empty vector (pYTK-e112) or a plasmid expressing Doa10 (either wild-type or Δ843-883 [“ΔL8/9b”]) under a *GAL1* promoter.

Overnight culture was diluted to an OD_600_ of 0.2 in 10 ml of fresh minimal medium SC(−Leu/−Ura) containing 2% raffinose, grown at 30°C until OD_600_ reached 0.5, and induced with 2% galactose for 4 h. Cells were harvested, washed with deionized water, and resuspended in 200 µL buffer LB with protease inhibitors. Cells were then lysed by bead-beating. After removing the glass beads, the cell lysate was supplemented with 1% LMNG and 0.2% CHS and incubated at 4°C for 1 h. The lysate was then clarified by centrifugation and incubated with Sepharose resins conjugated with GFP nanobody for 3 h. The resins were washed three times with buffer WB and proteins bound to the resins were eluted by SDS sample buffer.

### SDS-PAGE and antibodies (KW)

To measure the expression level of Doa10, 2.0 OD_600_ units of cells were collected, resuspended in 230 µl of 0.26 M NaOH and 0.13 M β-mercaptoethanol. Cell suspensions were incubated at room temperature for 5 min, then spun down by centrifugation at 6000 *g* for 3 min. Pellets were resuspended with 50 µl of reduced SDS sample buffer. Cell lysates were incubated at 37°C for 30 min and clarified by centrifugation at 21,000 *g* for 10 min before analysis by SDS-PAGE. SDS-PAGE was performed using Tris-glycine gels, except for Fig.5b,c where 4-12% Bis-Tris SDS-PAGE gels (Thermo Fisher) were used.

Immunoblotting experiments were performed with anti-Doa10 antiserum (a gift from M. Hochstrasser; 1:1,000 dilution), anti-GFP antibody (Thermo Fisher #MA5-15256 ; 1:3,000 dilution), anti-ALFA-tag nanobodies fused with a rabbit Fc domain (home-made), anti-Strep-tag antibody (Genscript #A01732, ; 1:2,000 dilution), anti-FLAG-tag antibody (Sigma #F1804; 1,1000 dilution), anti-Pgk1 antiserum (a gift from J. Thorner; 1:1,000 dilution). Secondary antibodies used in this study were goat anti-rabbit (Thermo Fisher #31460; 1:10,000 dilution), goat anti-mouse (Thermo Fisher #31430; 1:10,000 dilution).

## Acknowledgements

We are indebted to M. Hochstrasser for yeast strains, plasmids, the Doa10 antiserum, and helpful discussions. We thank J. Thorner for the Pgk1 antiserum. This work was supported by the Vallee Scholars Program (E.P.), Pew Biomedical Scholars Program (E.P.), Hellman Fellowship (E.P.), Jane Coffin Childs postdoc fellowship (K.W.). J.C.G. acknowledges support from NIH (R01 GM123169). Computational resources were provided through XSEDE (TG-MCB130173), which is supported by NSF (ACI-1548562). This work also used the Hive cluster, which is supported by the NSF (1828187) and is managed by PACE at Georgia Tech.

## Authors contributions

K.W., performed biochemical experiments, except for experiments in Fig. 6 and Supplementary Fig. 8e, which were performed by Y.C. S.I. purified samples for cryo-EM analysis and performed initial biochemical studies. E.P. and S.I. collected and analyzed cryo-EM data, and built atomic models. E.P. performed AlphaFold2 modeling. D.L. and J.C.G. performed MD simulations and analyses. A.T. assisted K.W. on biochemical experiments. E.P. supervised the project and wrote the manuscript with input from all authors. All authors contributed to data interpretations and manuscript editing.

## Competing interests

Authors declare no competing interests.

## Supplementary information

Supplementary Information is available for this paper.

## Corresponding authors

Correspondence and requests for materials should be addressed to Eunyong Park

## Data availability

The cryo-EM map and model are available through EM Data Bank (EMDB) and Protein Data Bank (PDB) under the following accession codes, respectively: EMD-41508 and PDB ID 8TQM.

## Notes

### Competing Interest Statement

The authors have declared no competing interest.

