## Supplementary Figures and Tables for "Substrate recognition mechanism of the endoplasmic reticulum-associated ubiquitin ligase Doa10"

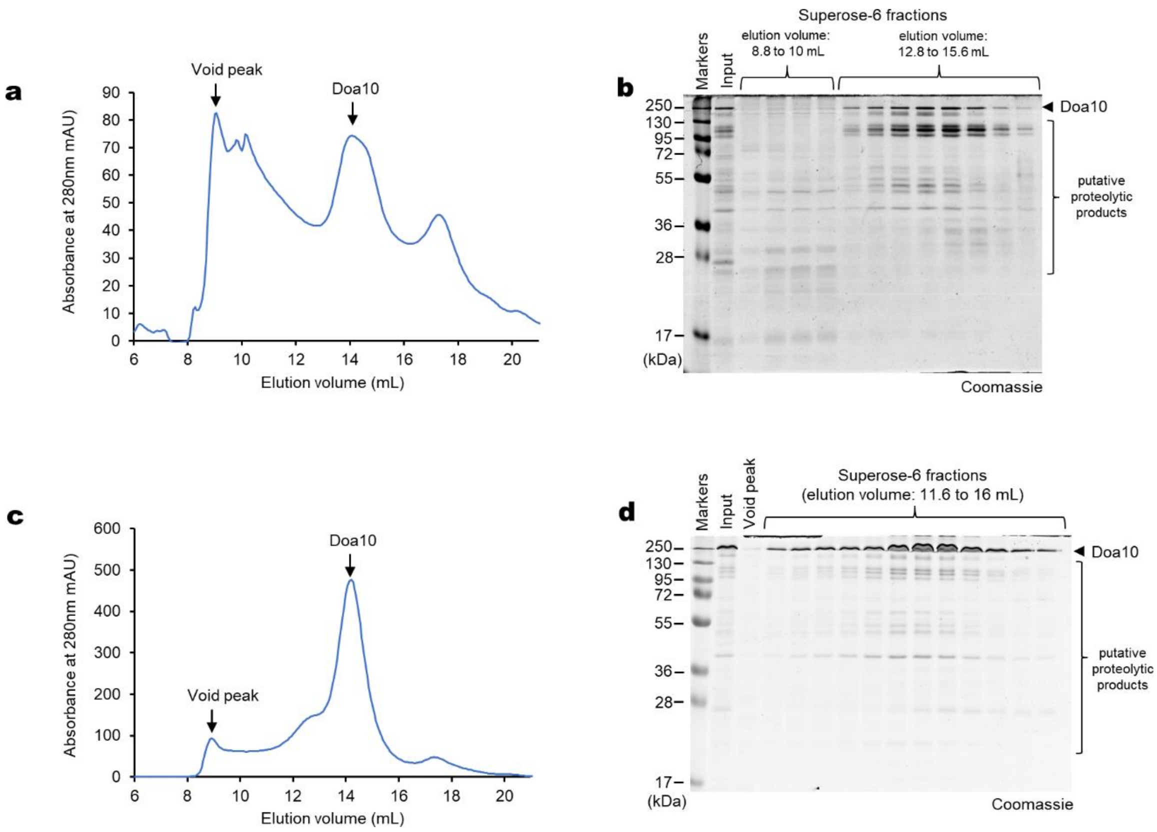

**Supplementary Figure 1. Purification of Doa10 from *S. cerevisiae*.**
**a**, Superose-6-increase size-exclusion chromatography (SEC) elution profile of affinity-purified endogenous Doa10 (GFP-tagged Doa10). **b**, Coomassie-stained SDS gel of Superose 6 fractions shown in **a**. **c** and **d**, As in **a** and **b**, but Doa10 was overexpressed by replacing the endogenous promoter with a *GAL1* promoter.

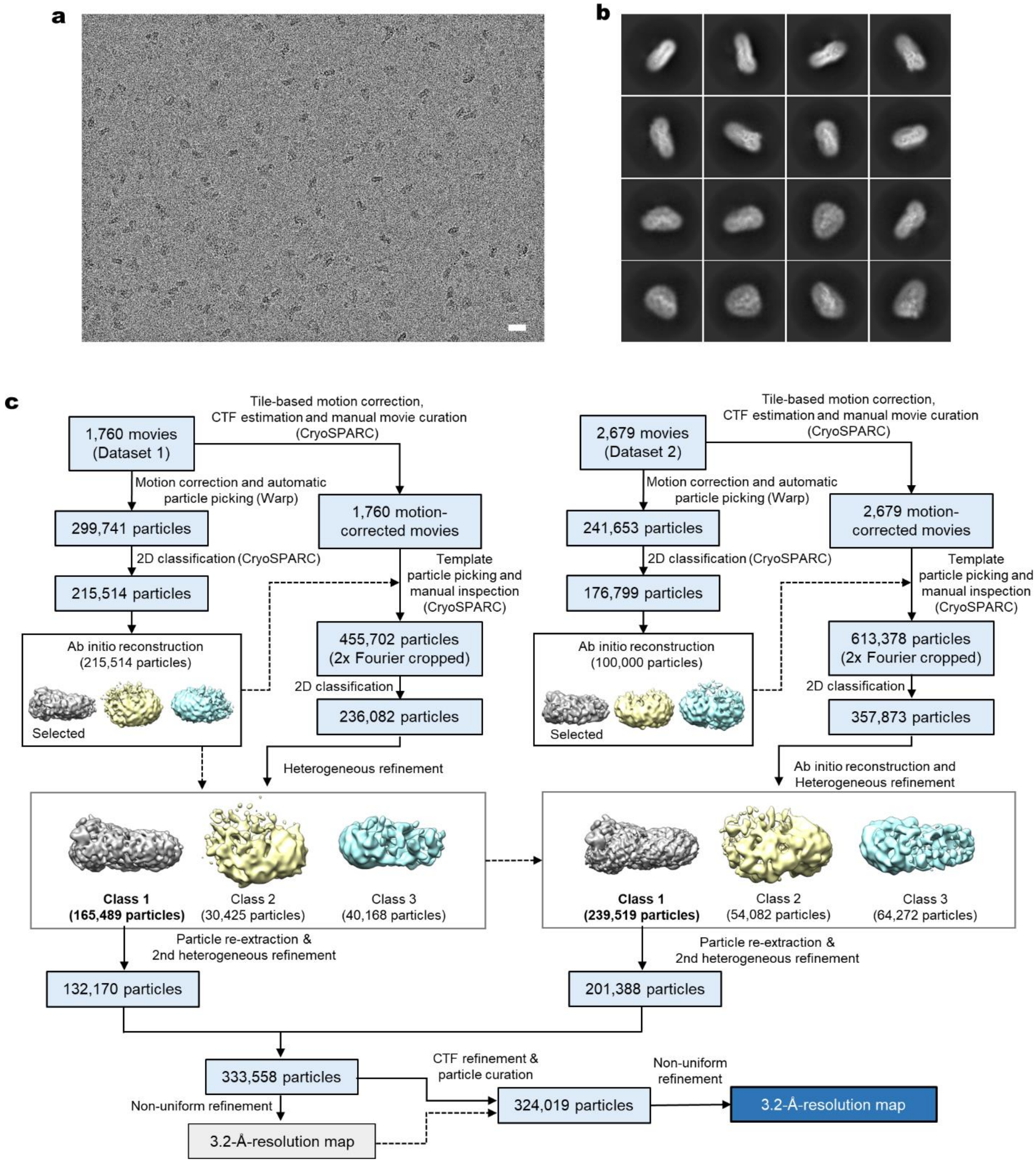

**Supplementary Figure 2. Cryo-EM analysis of Doa10.**
**a**, A representative micrograph (cropped) image of Doa10 particles. Scale bar, 200 Å. **b**, Representative 2D class averages. **c**, Schematic diagram for the cryo-EM image analysis procedure.

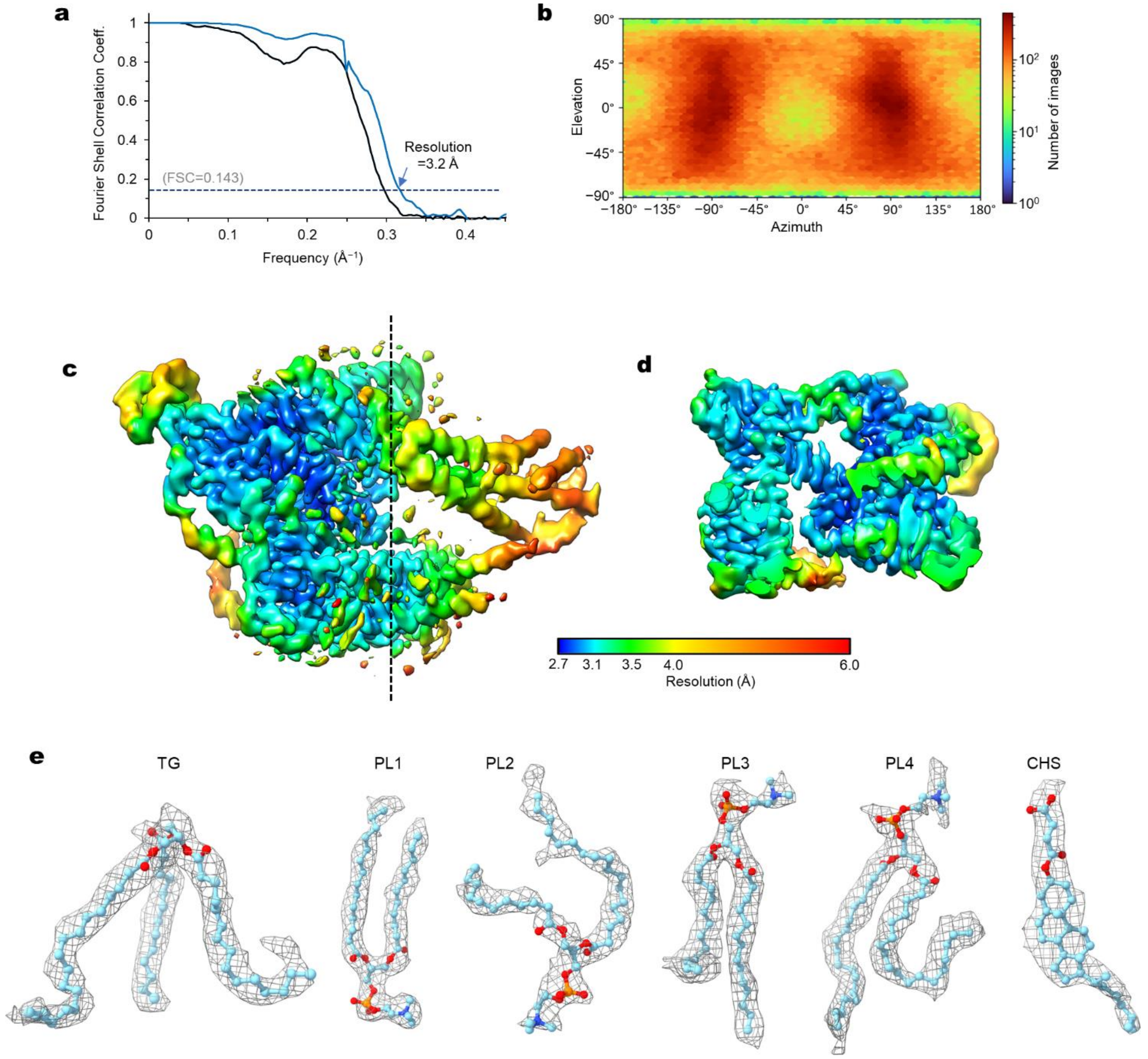

**Supplementary Figure 3. Quality of the cryo-EM map of Doa10.**
**a**, Fourier shell correlation (FSC) between the two half maps of the final 3D reconstruction. Blue, tight mask and corrected for masking; black, spherical mask. **b**, Distribution of particle orientation. **c** and **d**, Local resolution map (isosurface is an unsharpened map). The dash line indicates the cutaway plane for the view shown in **d**. **e**, Densities (gray mesh) of ordered lipids identified in the Doa10 cryo-EM structure.

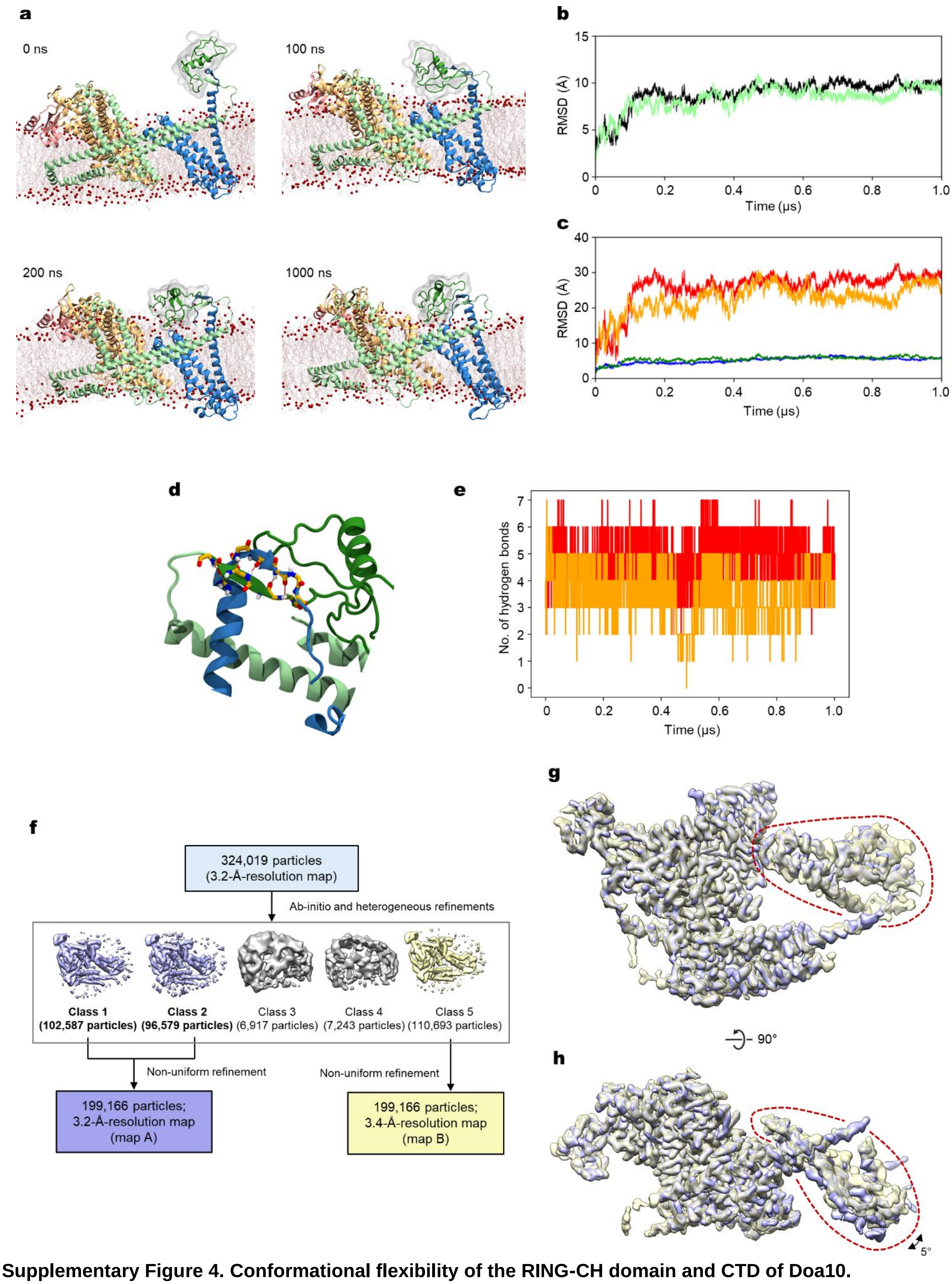

**Supplementary Figure 4. Conformational flexibility of the RING-CH domain and CTD of Doa10.**
(see next page for legend)

**Supplementary Figure 4. Conformational flexibility of the RING-CH domain and CTD of Doa10.**
**a**, Snapshots of MD simulations of wild-type Doa10 in a model membrane. The RING-CH domain is shown as a green ribbon and gray semi-transparent surface. **b**, Root-mean-square deviation (RMSD) using the Cα atoms for the two replicas of the Doa10 MD simulations. **c**, As in **b**, but additionally showing RMSD calculated for the RING-CH domain (red and orange) or the transmembrane domain (blue and green) for each replica. **d**, Snapshot of the MD simulations highlighting the two-strand β-sheet formed between the RING-CH and CTE of Doa10. **e**, The number of hydrogen bonds between the RING-CH and CTE over the duration of the MD simulations. Red and orange traces represent the two independent replicas. **f**, Additional classification of the Doa10 cryo-EM dataset to analyze the conformational heterogeneity of CTD. **g** and **h**, The two maps obtained from the procedure in **f** are overlaid. Shown are top (cytosolic; panel **g**) and side (panel **h**) views. CTD is indicated by red dashed lines.

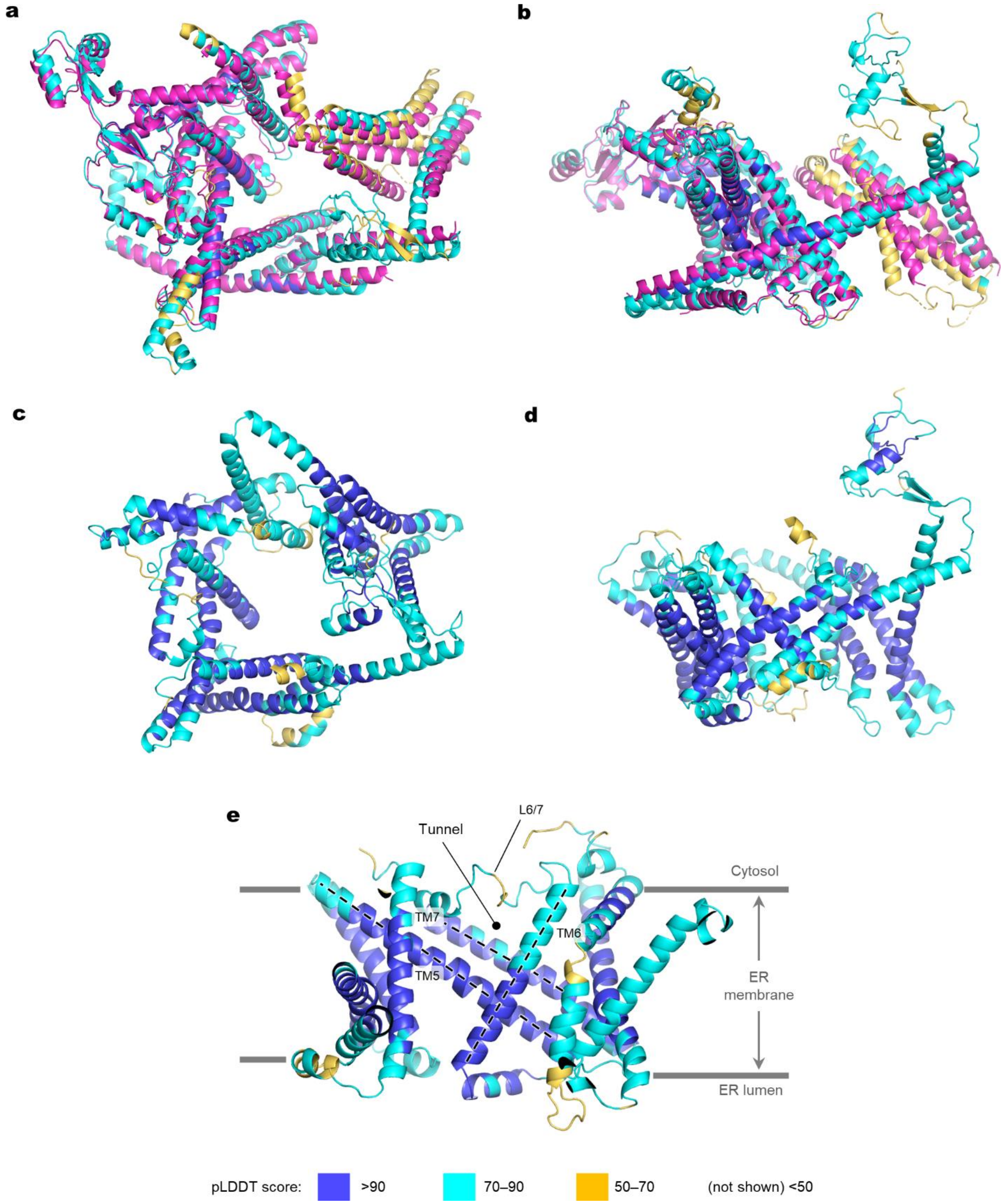

**Supplementary Figure 5. AlphaFold2 models of yeast Doa10 and human MARCH6.**
**a** and **b**, Comparison between the cryo-EM structure (magenta) and AlphaFold2 model (other colors) of yeast Doa10. The AlphaFold2 model is colored according to the pLDDT score (blue, >90; cyan, 70 to 90; yellow, 50 to 70). Parts below less than a pLDDT score below 50 were not shown. Shown are top (cytosolic; panel **a**) and side (panel **b**) views. **c**–**e**, AlphaFold2 model of human MARCH6. Top (**c**) and side (**d**) views, and a view into the lateral tunnel from the central cavity (**e**).

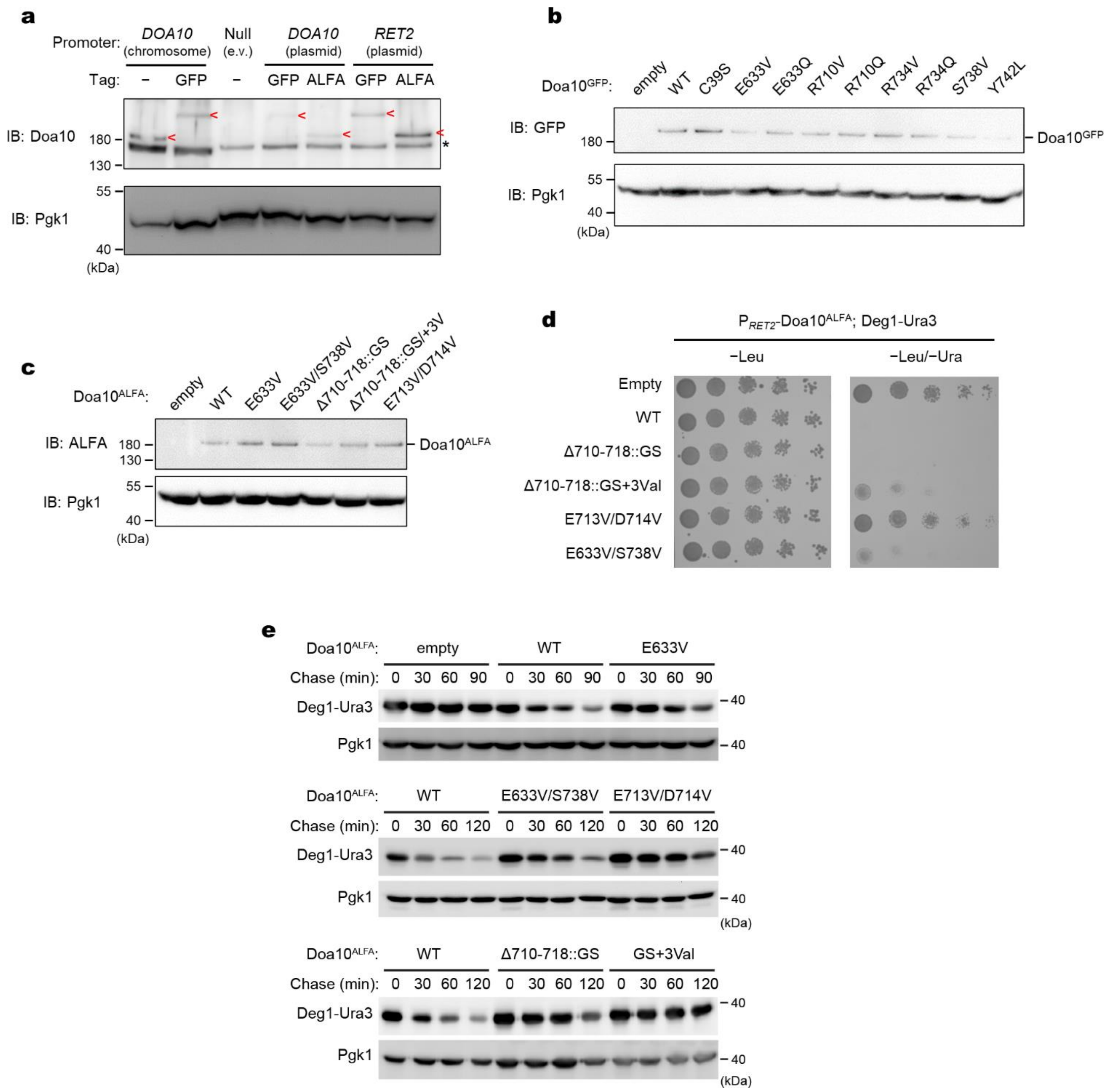

**Supplementary Figure 6. Effects of Doa10 mutations on Deg1-Ura3 degradation.**
**a,** Comparison of the endogenous and exogenous expression levels of Doa10 with either a C-terminal GFP-tag or ALFA-tag. Protein levels were measured by anti-Doa10 immunoblotting. 3-phosphoglycerate kinase (Pgk1) was used for a loading control. **b** and **c**, Expression levels of WT Doa10 and indicated mutants with a C-terminal GFP-tag (panel b) or with a C-terminal ALFA tag (panel c) was measured by immunoblotting. All the variants were expressed from a Doa10 promoter in a CEN/ARS plasmid. Pgk1 was used for a loading control. **d**, Yeast growth inhibition assay was performed as in Fig. 4e, but expressing Doa10 under the *RET2* promoter. Note that a higher expression level of Doa10 under the *RET2* promoter compared to the one under the endogenous Doa10 promoter (panel a) produces an overall stronger growth inhibition. **e**, Cycloheximide chase analysis of Deg1-Ura3 with indicated Doa10 mutants. Deg1-Ura3 levels were detected by anti-Strep-tag immunoblotting. Pgk1 was used for a loading control. See Fig. 4f for mean±s.e.m. from three independent experiments.

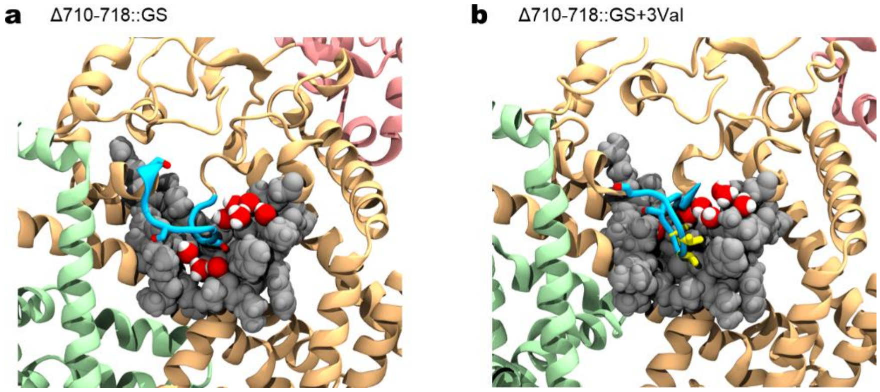

**Supplementary Figure 7. Example snapshots of MD simulations on Doa10 L6/7 mutants.** As in Fig. 4 g–i, but showing the Δ710-718::GS and Δ710-718::GS+3Val mutants of Doa10.

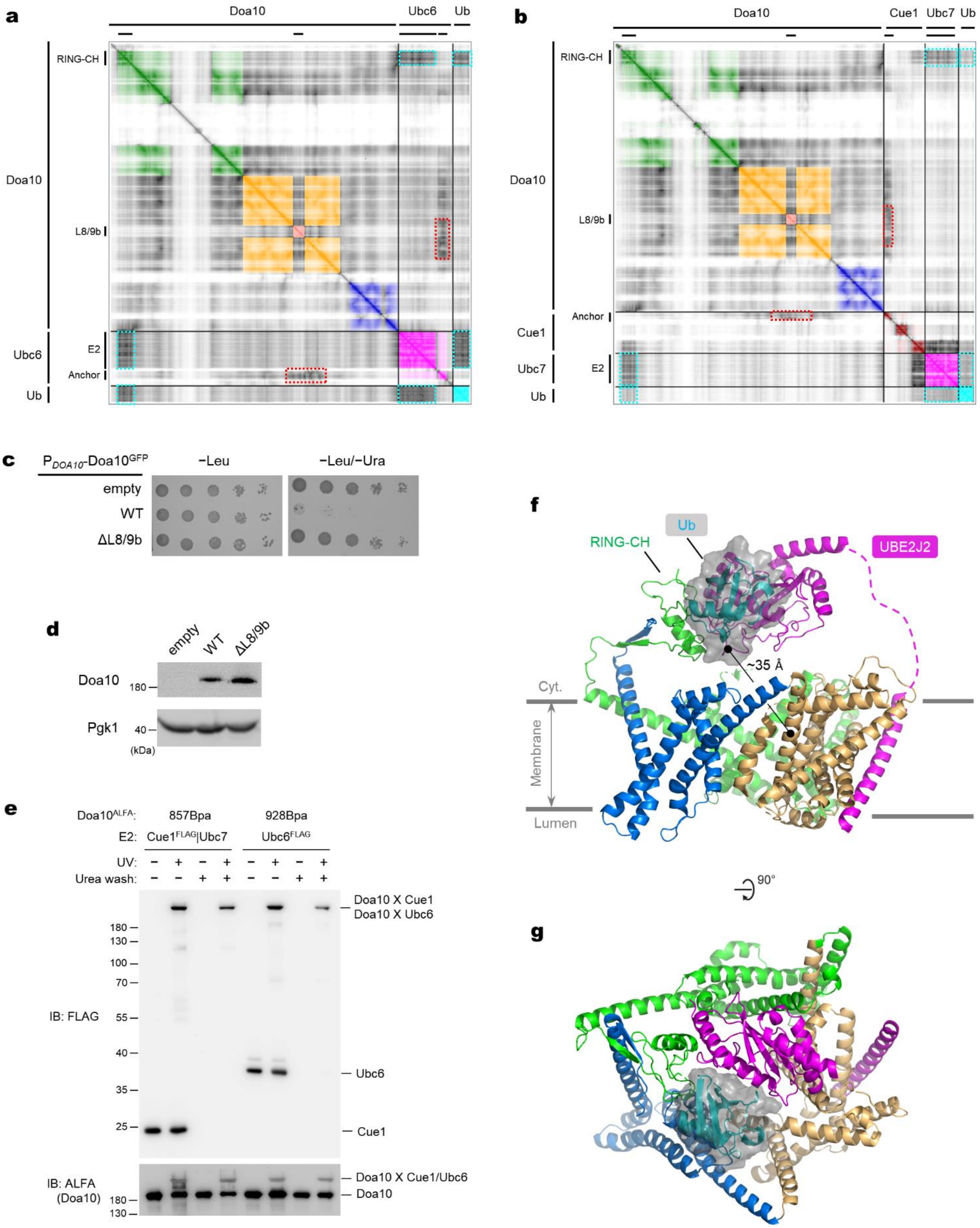

**Supplementary Figure 8. Interactions between Doa10 and E2s.**
(see next page for legend)

**Supplementary Figure 8. Interactions between Doa10 and E2s.**
**a**, Predicted Aligned Error (PAE) matrix for the AlphaFold2 model of the Doa10–Ubc6–Ub complex. Regions are colored according to the domain color scheme in Fig. 6 (only regions with >50 pLDDT score are in color). Dashed boxes in cyan indicate the regions corresponding to contacts between the RING-CH, E2 domain, and Ub. Dashed boxes in red indicate the contacts between the TM anchor of Ubc6 and Doa10. **b**, As in **a**, but for AlphaFold2 model for Doa10–Cue1–Ubc7–Ub. **c**, Yeast growth inhibition assay comparing the activities of WT and ΔL8/9b Doa10. Doa10 with a C-terminal GFP-tag were expressed from a *DOA10* promoter in a CEN/ARS plasmid. **d**, Expression levels of WT and ΔL8/9b Doa10 were measured by anti-GFP immunoblotting. Pgk1 was used as a loading control. **e**, Verification of covalent, UV-dependent crosslink adducts. Where indicated, ALFA nanobody beads were additionally washed with 6 M urea during immunoprecipitation to dissociate non-covalently associated proteins. **f** and **g**, As in Fig. 6 a and b, but showing an AlphaFold2 model of a complex of human MARCH6, UBE2J2, and Ub. Panels in **f** and **g** show side and cytosolic views, respectively.

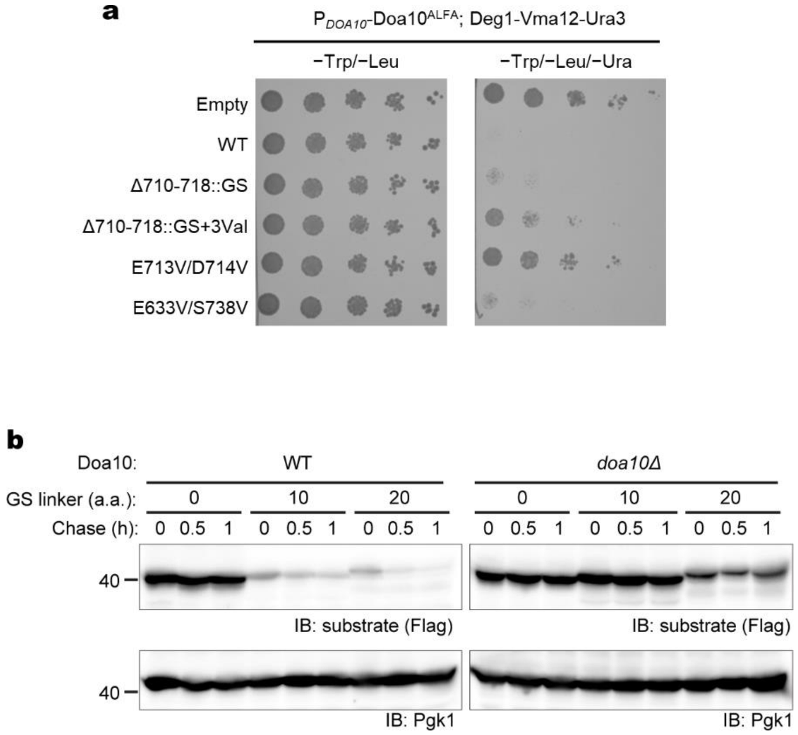

**Supplementary Figure 9. Doa10-dependent degradation of Deg1-Vma12-Ura3.**
**a**, Yeast growth inhibition assay with Deg1-Vma12-Ura3 and indicated Doa10 mutants. Deg1-Vma12-Ura3 was expressed under the *MET25* promoter, and all Doa10 mutants were expressed from the *DOA10* promoter. **b**, Cycloheximide chase analysis of Deg1$_{1\text{-}35}$-Vma12$_{\Delta132}$-Ura3 with different lengths of the GS linker between Deg1$_{1\text{-}35}$ and Vma12$_{\Delta132}$. Deg1 substrates were detected by anti-FLAG-tag immunoblotting (a FLAG-tag is attached to the C-terminus of Ura3). Pgk1 was used for loading control. See Fig. 7d for mean±s.e.m. from three independent experiments.

**Table S1. Cryo-EM data collection, refinement and validation statistics.**

| | **Doa10 from *S. cerevisiae*** (PDB:8TQM, EMDB: 41508) |
|---|---|
| **Data collection and processing** | |
| Microscope | FEI Titan Krios G2 |
| Magnification | 64,000x |
| Voltage (kV) | 300 |
| Electron exposure ($e^{-}/Å^2$) | 50 |
| Defocus range (μm) | −0.7 to −2.4 |
| Pixel size (Å) | 0.91 |
| Symmetry imposed | *C*1 |
| Initial particle images (no.) | 541,394 |
| Final particle images (no.) | 324,019 |
| Map resolution (Å) | 3.2 |
| FSC threshold | 0.143 |
| Map resolution range (Å) | 2.72 (highest), 7.15 (75% percentile) |
| **Refinement** | |
| Initial model used | De novo + AlphaFold2 model |
| Model resolution (Å) | 3.3 |
| FSC threshold | 0.5 |
| Map sharpening *B* factor ($Å^2$) | 112 |
| Model composition | |
| Non-hydrogen atoms | 7,231 |
| Protein residues | 843 |
| Ligands | 7 |
| *B* factors ($Å^2$) | |
| Protein | 34.10 |
| Ligand | 12.05 |
| R.m.s. deviations | |
| Bond lengths (Å) | 0.005 |
| Bond angles (°) | 1.055 |
| **Validation** | |
| MolProbity score | 1.70 |
| Clashscore | 12.0 |
| Poor rotamers (%) | 0 |
| Ramachandran plot | |
| Favored (%) | 97.47 |
| Allowed (%) | 2.53 |
| Disallowed (%) | 0 |
| CaBLAM outliers (%) | 0.73 |

**Table S2. List of yeast strains**

| Name | Genotype | Reference |
|---|---|---|
| MHY4086 | *MATa his3-Δ200 ura3-52 trp1-Δ63 leu2-3112 lys2-801::LYS2::*Deg1-Ura3 *doa10Δ::HphMX4* | Stuerner et al., 2012 |
| JY103 | *MAT***a** *his3-Δ200 ura3-52 trp1-Δ63 leu2-Δ1 lys2-801 ADE2* | Laney et al., 2006 |
| MHY10818 | *MATα his3-Δ200 leu2-Δ1 ura3-52 lys2-801 trp1-Δ63 doa10Δ::hphMX4* | Mehrtash et al., 2022 |
| BY4741 | *MATa his3-1, leu2-0, met15-0, ura3-0* | Horizon Discovery |
| ySI-118 | BY4741 *DOA10*-TEV-GFP::*natMX* | This study |
| ySI-154 | BY4741 *leu2*::$P_{GAL1}$-Doa10-TEV-GFP::*natMX* | This study |
| ySI-167 | BY4741 *doa10Δ*::*natMX* | This study |
| ySI-266 | MHY10818 *cue1Δ*::*natMX* | This study |
| yKW-283 | ySI-167 *leu2*::$P_{PGK1}$-Deg1-Ura3-2xstrep::*hphMX4* | This study |
| yYC-307 | MHY10818 *ubc6Δ*::*natMX* | This study |

**Table S3. List of plasmids**

| Name | Description | Reference |
|---|---|---|
| pYTK001 to 096 | Original MoClo YTK parts | Lee et al., 2015 |
| pYTK-e102 | *LEU2* integration vector containing a hygromycin marker (assembled from pYTK008, pYTK047, pYTK073, pYTK079, pYTK087, pYTK090, and pYTK093) | This study |
| pYTK-e205 | MoClo YTK part (type 4a) for 2x-Strep (Amino acid sequence: SGWSHPQFEKGGGSGGGSGGSAWSHPQFEK*) | This study |
| pYTK-e111 | CEN/ARS plasmid containing a Ura3 marker (assembled from pYTK008, pYTK047, pYTK073, pYTK074, pYTK081, and pYTK084 ) | This study |
| pYTK-e112 | CEN/ARS plasmid containing a Leu2 marker (assembled from pYTK008, pYTK047, pYTK073, pYTK075, pYTK081, and pYTK084) | This study |
| pDoa10-Doa10-split1 | Split pYTK-e112 (CEN/ARS)-$P_{DOA10}$-Doa10(1-608) | This study |
| pRET2-Doa10-split1 | Split pYTK-e112 (CEN/ARS)-$P_{RET2}$-Doa10(1-608) | This study |
| pTDH3-Doa10-split1 | Split pYTK-e112 (CEN/ARS)-$P_{TDH3}$-Doa10(1-608) | This study |
| Doa10–GFP-split2 | Split pYTK-e112 (Leu2 marker)-Doa10(608-1319)-TEV-GFP (Amino acid sequence of the tag: GTGSGTGENLYFQGTASGGGSKGEELFTGVVPILVELDGDVNG HKFSVSGEGEGDATYGKLTLKFICTTGKLPVPWPTLVTTFGYGV QCFARYPDHMKQHDFFKSAMPEGYVQERTIFFKDDGNYKTRAE VKFEGDTLVNRIELKGIDFKEDGNILGHKLEYNYNSHNVYIMADK QKNGIKVNFKIRHNIEDGSVQLADHYQQNTPIGDGPVLLPDNHYL STQSALSKDPNEKRDHMVLLEFVTAAGITHGMDELYKVDLDK*) | This study |
| Doa10-ALFA-split2 | Split pYTK-e112 (Leu2 marker)-Doa10(608-1319)-ALFA (Amino acid sequence of the tag: GTSRLEEELRRRLTE*) | This study |
| pKW043 | MoClo YTK part (type 3a) for Deg1 | This study |
| pKW050 | MoClo YTK part (type 3b) for Ura3 | This study |
| pKW155 | pYTK-e102-PGK1-Deg1-Ura3-2xstrep. Assembled from pYTK-e102, pYTK011 ($P_{PGK1}$), pKW043 (pYTK001-Deg1), pKW050 (pYTK001-Ura3), pYTK001-e205 (2xStrep), pYTK061 (tENO1) | This study |
| p414-Deg1-Vma12-Ura3 | Plasmid expressing Deg1-Vma12-Ura3 from a *MET25* promoter | Ravid et al. 2006 |
| pSK-B399-GFP-NAT | Template plasmid for PCR to generate a DNA fragment containing a C-terminal TEV-GFP tag and natMX (for chromosomal tagging). | Gift from the S. Klinge lab |
| pYC-300 | pYTK-e111-$P_{GAL1}$-Cue1-2xFLAG :: $P_{GAL1}$-Ubc7-2xStrep | This study |
| pYC-301 | pYTK-e111-$P_{GAL1}$-3xFLAG-6xHis-Ubc6 | This study |
| pYC-302 | pYTK-e111-$P_{GAL1}$-Cue1-2xFLAG :: $P_{GAL1}$-Ubc7-2xSPOT :: $P_{GAL1}$-3xFLAG-6xHis-Ubc6 | This study |

**Supplementary Movie 1. MD simulation on the cryo-EM/AlphaFold2 hybrid model of Doa10.**
The hybrid model was placed in a model lipid bilayer, and an all-atom MD simulation was performed for a 1-µs duration. Shown is the side view along the membrane plane with Doa10 in a ribbon representation and positions of lipid headgroups indicated by red spheres. The color scheme is the same as in Fig. 1. The hybrid model includes the N-terminal RING-CH domain, for which an additional surface representation (surface in transparent gray) is used.

**Supplementary Movie 2. Zoom-in views into the lateral tunnel of WT and mutant Doa10 in MD simulations.**
Parts of TMs 5 to 7 that form the lateral tunnel are shown (front view). In the WT, E633V/S73V, and E713V/D714V panels, the L6/7 loop (positions 710-718) is shown in cyan licorice, whereas in the Δ710-718::GS (‘GS’) and Δ710-718::GS+3Val (‘GS+3Val’) mutants, the equivalent segment is shown in blue and green licorice, respectively. Gray spheres are used for a space-filling representation of the wedge-shaped tunnel-lining amino acids from TMs 5 to 7. Yellow residues are mutated valine.
